# Y chromosome shredding in *Anopheles gambiae*: insight into the cellular dynamics of a novel synthetic sex ratio distorter

**DOI:** 10.1101/2024.05.09.593338

**Authors:** Matteo Vitale, Nace Kranjc, Jessica Leigh, Kyrous Kyrou, Thomas Courty, Louise Marston, Silvia Grilli, Andrea Crisanti, Federica Bernardini

## Abstract

Despite efforts to explore the genome of the malaria vector *Anopheles gambiae*, the Y chromosome of this species remains enigmatic. The large number of repetitive and heterochromatic DNA sequences makes the Y chromosome exceptionally difficult to fully assemble, hampering the progress of gene editing techniques and functional studies for this chromosome. In this study, we made use of a bioinformatic platform to identify Y-specific repetitive DNA sequences that served as a target site for a CRISPR/Cas9 system. The activity of Cas9 in the reproductive organs of males caused damage to Y-bearing sperm without affecting their fertility, leading to a strong female bias in the progeny. Cytological investigation allowed us to identify meiotic defects and investigate sperm selection in this new synthetic sex ratio distorter system. In addition, alternative promoters enable us to target the Y chromosome in specific tissues and developmental stages of male mosquitoes, enabling studies that shed light on the role of this chromosome in male gametogenesis. This work paves the way for further insight into the poorly characterised Y chromosome of *Anopheles gambiae*. Moreover, the sex distorter strain we have generated promises to be a valuable tool for the advancement of studies in the field of developmental biology, with the potential to support the progress of genetic strategies aimed at controlling malaria mosquitoes and other pest species.

**Author summary:** Genetic elements known as sex ratio meiotic drive can manipulate the sex ratio of offspring, favouring the male or female sex. This fascinating phenomenon has inspired the development of synthetic sex ratio distorter systems in several organisms. In species where females and males harbour XX and XY sex chromosomes respectively, the X-chromosome can be ‘shredded’ during male gametogenesis, leading to the production of non-functional X-bearing sperm, while Y-bearing sperm are left intact and able to fertilise the eggs. These systems can produce offspring that are extremely biased towards males, which can be used as genetic tools to control harmful insect populations. In our study, we applied this molecular strategy to target the Y chromosome of *Anopheles gambiae*. Our aim was to investigate the cellular consequences of the shredding of this chromosome, the impact on meiosis and sperm selection, and the potential to achieve strong female bias in the offspring. The outcome of this study enhances our understanding of the molecular and biological mechanisms behind synthetic sex-ratio distorters in Anopheles mosquitoes, which could inform the development of vector control strategies that target sex ratio. Additionally, we present a genetic sexing strain able to produce mostly females, providing a valuable genetic tool for fundamental studies on this deadly vector.

## Introduction

In unisexual organisms, sexual development commonly depends on sex chromosomes, which can be classified as homomorphic or heteromorphic. While homomorphic sex chromosomes exhibit only a few differences and are almost indistinguishable, heteromorphic sex chromosomes can drastically differ in their size and shape (1–3).

In most cases, heteromorphic sex chromosomes are thought to derive from a pair of ancestral autosomes that undertook a series of unique evolutionary processes (4–6). In the case of XY sex determination systems, the classic model assumes that these processes are initiated by the acquisition of a sex-determining factor on a proto-Y chromosome. Subsequently, due to the accumulation of sexually antagonistic mutations, recombination is suppressed in the chromosomal region involved in sex determination (5,7). In addition, genetic phenomena such as Muller’s rachet and genetic hitchhiking would contribute to progressive degeneration, loss of gene content, and accumulation of repetitive DNA (4,5). In a few rodent species, the degeneration process is exacerbated to the extent that the Y chromosome is lost entirely (8,9). The scarcity of protein-coding genes and high prevalence of repetitive DNA make the Y chromosome a ‘genetic desert’, mostly heterochromatic, while the X chromosome usually retains most of its ancestral euchromatic state (2,5,6,10). The loss-of-function of Y-linked genes creates an unbalance in the gene expression level in the heterogametic sex, which is re-established through the evolution of dosage compensation mechanisms (10–13).

Male mosquitoes belonging to the *Anopheles* complex, have heteromorphic sex chromosomes that, with the exception of *An. merus*, are karyotypically distinct (14–16). A number of genomic and functional studies have been conducted on sex chromosomes and sex determination in the main malaria vector *Anopheles gambiae* (17–25). This species has a pair of fully differentiated heteromorphic X and Y chromosomes with some level of sequence similarity, mostly involving satellite DNA. In addition, some evidence suggests that recombination between these chromosomes can still occur in limited regions (14,26).

The Y chromosome carries the male determining gene, *yob,* along with seven other genes with paralogues on the autosomes and X chromosome. The function of these genes has not been fully characterised but RNA sequencing data have shown expression, for some of these genes, along several developmental stages and in male reproductive tissues (14,18,21). Although a full chromosomal assembly for this chromosome is not available, experiments based on long-read sequencing in conjunction with cytogenetic mapping suggest that most of its content, around 92.3 %, is represented by transposable elements (TEs) and satellite DNA (stDNA) (14,27). TEs are actively expressed throughout mosquito development and in reproductive tissues, while the most abundant stDNA families, AgY477 and AgY373, are only expressed in the male germline (14).

High degrees of dynamism in Y-linked repetitive sequences have been shown among different species of the Anopheles complex (14,15). Despite these differences, empirical studies have demonstrated the possibility, within the Anopheles complex, of inter-species Y chromosome introgression, ruling out the role of this chromosome in hybrid incompatibility (17). For some species of the *Anopheles* genus, evidence has pointed to the involvement of the Y chromosome in mating behaviour (28).

In recent years, a number of genetic vector control strategies have been developed to target *Anopheles* mosquitoes (29–38). Among these strategies, synthetic sex ratio distortion (SSD) systems have gained great attention (32,33,39–41). These systems rely on manipulating the sex ratio of the offspring, favouring males, to suppress or eliminate a target population. Existing SSD systems targeting *An. gambiae* are based on the use of endonucleases, I-Ppol or Cas9, that cleave the ribosomal DNA repeats (rDNA) located, in this species, exclusively on the X chromosome (32,33). The chromosomal damage induced by the endonuclease activity, expressed during male gametogenesis, results in the generation of non-viable X-bearing sperm, while Y-bearing sperm are left viable and able to fertilise the eggs. A cytological investigation of the *An. gambiae* SSD strain ^gfp^124L-2, based on the activity of the endonuclease I-Ppol and referred to herein as Ag(PMB)1, revealed that, upon mating, both X- and Y-bearing sperm are transferred to female spermathecae in the same ratio, suggesting that selection of functional Y-bearing sperm is operated in the female through mechanisms that remain elusive (42).

Recently, a pest control strategy based on Y chromosome shredding coupled with a homing mechanism has been investigated in a mouse model (43). Stochastic modelling has indicated that eradication of pest vertebrate populations can be achieved by the depletion of XY males, thereby introducing a strong female-bias into the population (43). However, while modelling has shown promising figures, efficient Y-chromosome elimination has only been demonstrated in cultured cells and early-stage embryos (44,45).

In this study, we used CRISPR/Cas9-based nucleases to target Y-chromosome specific sequences and to develop the first SSD system causing female bias in *An. gambiae*. Although, a vector control strategy based on biassing the progeny towards females would not be desirable for our target species, given that malaria is transmitted by the bite of infected female mosquitoes, the bioinformatic approach and the cytologic observations presented in this study could be valuable to explore the potential of such SSD system in other pest species where the female sex is not directly causing harm and where mathematical modelling shows promising prospects for population control.

The strain we have generated represents a valuable tool for genetic sexing in mosquitoes and offers a unique opportunity to carry out functional studies and investigate the biology of the Y chromosome. Our results significantly increase our knowledge of fundamental aspects of *An. gambiae* male gametogenesis and reproduction. Moreover, this work sheds light on the cytological mechanisms underlying SSD systems and sex chromosome behaviour. This knowledge has the potential to facilitate the development of novel genetic technologies based on this strategy.

## Results

### 1 Identification of Y chromosome-specific sequences for the development of a Y-shredder system in *Anopheles gambiae*

To identify CRISPR/Cas9 target sequences in the *An. gambiae* Y chromosome, we used the assembly-free computational pipeline Redkmer (46). This bioinformatic pipeline was originally designed for the identification of unique and highly repetitive X chromosome-linked sequences that could serve as target sites for X chromosome shredding systems (47,48). For the analysis, short-read whole genome sequencing (WGS) data obtained from both female and male individuals is used. These short-reads are aligned to long-read WGS datasets (Oxford Nanopore Technology, ONT) generated from both males and females. Chromosome Quotient (CQ) is calculated as the ratio of coverage of female *versus* male short-reads mapped to the long-reads. CQ is then used to assign the chromosomal origin of the long-reads to 3 bins: autosomal (CQ ∼1), X-specific (CQ ∼2), and Y-specific (CQ ∼0). Lastly, the long-read sequences from chromosomal bins are used as a reference to allocate 25 bp long k-mers originated from the previously mapped short-read sequences. Redkmer pipeline was modified to filter and identify Y chromosome-specific and highly repetitive k-mers that could be used as a target for the development of a CRISPR/Cas9-based shredding system (Figure 1A). The pipeline identified 1,795 candidate Y-specific k-mers, of which 866 showed no off-target in the autosomal and X chromosome bins. As CRISPR/Cas9 target sequences require 20 bps homology with a gRNA, among the candidates, we selected k-mers with a minimum sequence length of 20 bps containing a Protospacer Adjacent Motif (PAM) at either the 5’ or the 3’ end, leading to a final list of 175 candidate k-mers (Figure 1A). The alignment of the 1,795 Y-specific k-mers to the available *An. gambiae* reference genome (AgamP4) allowed us to map potential target sites *in silico*. 1,395 out of 1,795 k-mers corresponded to sequences of Y-linked satellite and retrotransposon, largely described in previous studies (14,27) (BAC60P19 and BAC10L10 are classified as *Zanzibar* retrotransposons) (Figure 1B).

**Figure 1:**
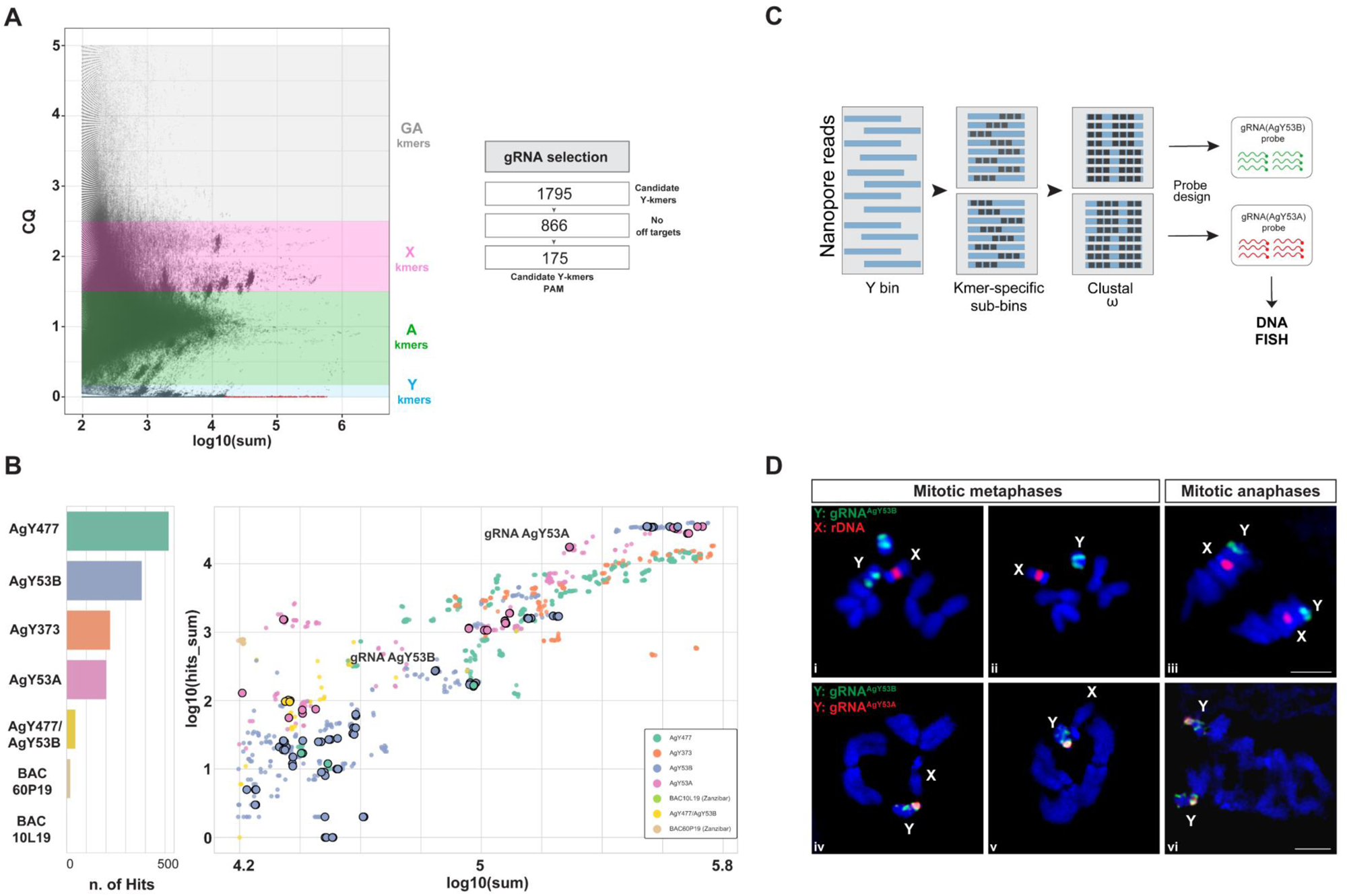
Identification and validation of Y chromosome-specific target sequences for the development of a CRISPR/Cas9-based Y shredder system. (A) On the left, a plot showing abundance (in log10 of sum, X-axis) *versus* Chromosome Quotient (CQ, Y-axis) of k-mers generated by the RedKmer pipeline. In red, predicted Y chromosome k-mers to be used as targets for CRISPR/Cas9 systems. On the right, the filtering steps (including off-target and PAM motif analyses) used to generate the final list of Y chromosome k-mer candidates. (B) On the left, *in silico* characterization of candidate Y-k-mers using BlastN showing matches with the *An. gambiae* genome reference (BAC 60P19 and 10L19 are classified as *Zanzibar RT*). On the right, the dot plot shows the coverage of the candidate Y-k-mers in the short-read dataset (X-axis, log10(sum)) and long-read dataset (Y-axis, log10(hits_sum)). The colour code is based on k-mer sequence identity following the mapping using BlastN. Candidate Y-k-mers (≤ 25 bp length) showing no-off targets in X or autosomal bins and PAM motif are circled (n=175). The final k-mers selected for empirical validation are highlighted with the names gRNA AgY53A and AgY53B. (C) Schematic representation of the workflow used to generate DNA FISH probes specific to gRNA AgY53A and AgY53B. Each k-mer sequence is aligned to the Y-bin long-reads using bowtie1 to extract k-mers-specific ONT reads and generate k-mers-specific sub-bins. Multiple sequence alignment software (Clustal Omega) is used to generate a consensus sequence from k-mers-specific ONT reads. The consensus sequence is used to design > 40 bps-long probes. Black boxes indicate tandem arrays of the satellite repeats. (D) DNA FISH performed on gonial cell mitotic chromosomes at different stages obtained from *An. gambiae* WT testis. Panels i-iii show complete Y-linkage of the probe generated from gRNA AgY53B (green) in association with a probe specific to the rDNA locus located on the X chromosome (red). Note that the additional green signal in panel i denotes a Y chromosome deriving from the genetic makeup of an adjacent cell. Panels iv-v-vi show the Y chromosome linkage of the probes generated from gRNA AgY53B (green) and AgY53A (red). The signal from the probe specific to the gRNA AgY53B occupies a larger region on the Y chromosome when compared to AgY53A. In addition, the fluorescent signal from this probe is localised at one end of the Y chromosome;. X: X chromosome; Y: Y chromosome; A: autosomes. Plot A was generated by the RedKmer Pipeline, and plot B was largely inspired by Papathanos and Windbichler, 2018 (46).

We focused our attention on two k-mers belonging to the satellite families AgY53A and AgY53B (from now on referred to as gRNA-AgY53A and gRNA-AgY53B). gRNA-AgY53A and gRNA-AgY53B were selected due to their high repetitiveness, absence of off-targets, and low CQ values, making them suitable candidates as Y-linked target sites.

To cytologically validate the Y chromosome specificity of gRNA-AgY53B and gRNA-AgY53A, we performed Fluorescent In Situ Hybridization (FISH). Long-read sequences containing the gRNA target sequences were aligned to obtain the consensus sequence for the generation of FISH DNA probes. We designed 44 bp long DNA probes containing the gRNA sequence generated by RedKmer and including an extra sequence of the satellite tandem repeat to ensure a good level of signal detection. (Figure 1C). These probes were used to label mitotic and meiotic chromosomes obtained from *An. gambiae* wild type (WT) testes. In addition, DNA probes specific for the X-linked rDNA locus were used as controls (14,15). Probes specific to the gRNA-AgY53B showed complete Y-linkage in all analysed chromosome preparations, while X-linkage was observed for the control probe labelling the ribosomal locus (Figure 1D-i-ii-iii). As shown in Figure 1D-iv-v-vi, when using probes for both AgY53A and AgY53B, the Y chromosome was specifically labelled, but the signals differed in their spatial distribution along the chromosome. The signal from the gRNA-AgY53A probe has a dotted-like shape and is located at the distal end of the Y chromosome, however, given the highly heterochromatic composition of the Y chromosome and the absence of its full-length chromosomal assembly, it is difficult to classify this region as telomeric or centromeric. The fluorescent signal from the gRNA-AgY53B probe partially overlaps with gRNA AgY53A but occupies a more extensive region of the Y chromosome (Figure 1D-iv-v-vi). In addition, a detailed 3D model of the spatial distribution of the target sites is shown in Supplementary S1 Video. Based on its widespread distribution along the chromosome, we hypothesised that gRNA-AgY53B could lead to higher chromosomal damage; hence, we selected this as the target site for *in vivo* testing of Y chromosome shredding.

### 2 Targeting the Y-linked satellite AgY53B during male gametogenesis causes strong female bias in the progeny of *Anopheles gambiae*

To build a Y chromosome shredding system in *An. gambiae*, we generated a germline transformation construct, CRISPR^AgY53B^, containing a Cas9 endonuclease gene under the control of the spermatogenesis-specific *β2*-tubulin promoter (32,33,49) (Figure 2A). Ubiquitous expression of the gRNA cassette targeting satellite AgY53B was driven by the U6 promoter (50). In addition, the genetic construct harboured a Red Fluorescent Protein (RFP) and *piggyBac* inverted terminal repeat sequences for transposon-mediated random integration (Figure 2A). We injected *An. gambiae* embryos with the CRISPR^AgY53B^ genetic construct and a helper plasmid as a source of transposase (30,51,52). From the embryos surviving to hatching, individuals transiently expressing the fluorescent marker were identified. These individuals were crossed with their WT counterparts, and from their progeny, two independent transgenic strains were isolated and established.

**Figure 2:**
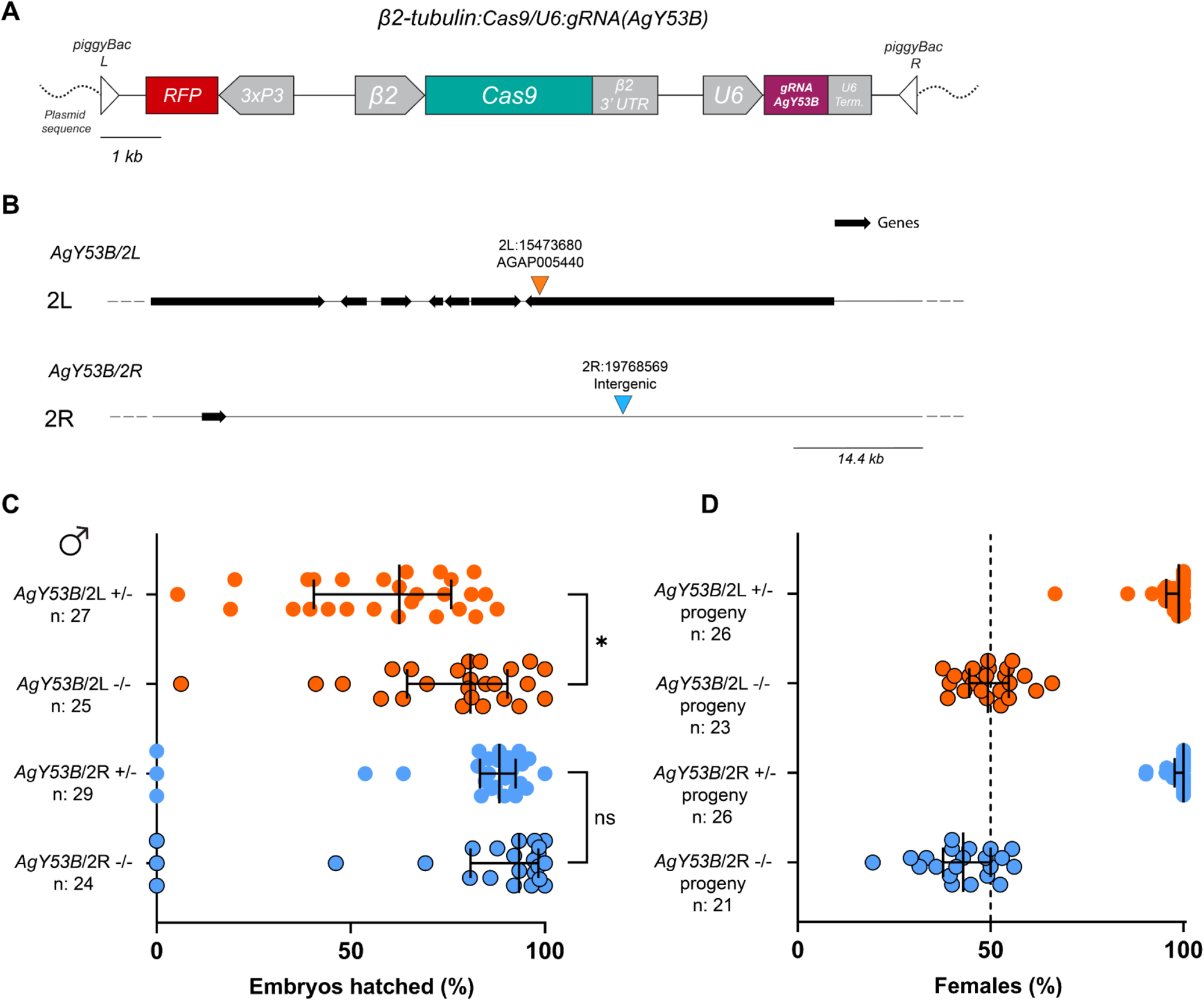
Generation of a Y chromosome shredding system in *An. gambiae*. (A) Schematic representation of the genetic construct used to target the Y-linked satellite AgY53B in *An. gambiae*. (B) Genome integration sites of the genetic construct in two independently generated transgenic strains, AgY53B/2L and AgY53B/2R. Dark arrows identify genes, while the black line indicates intergenic regions. (C) Phenotypic assay of AgY53B and control males. Strain AgY53B/2L +/- shows a significant reduction in the Hatching Rate (HR) compared to the control Agy53B/2L -/- (Dunn’s multiple comparison test (two-sided), *P value* = 0,0147). Strain AgY53B/2R +/- shows no reduction in HR when compared to the control AgY53B/2R -/- (Dunn’s multiple comparison test (two-sided), *P value* = 0,6984). Median and interquartile ranges are shown. (D) Percentage of females in the progeny of strains harbouring the CRISPR^AgY53B^ construct and control males. Progeny derived from AgY53B/2L +/- and Agy53B/2R +/- males exhibit a high level of sex ratio distortion towards females compared to the controls. The median of the percentage of females from the progenies is as follow: 94.28 %, 49.3 %, 100 %, 42.86 % (from top to bottom). Median and interquartile ranges are shown. Circled dots represent values from control samples. The dotted line in D shows the expected 50 % female ratio.

Molecular characterization revealed that in one of the transgenic strains, the genetic construct had integrated on chromosome arm 2L at position 15473680 within the 3’ UTR of the uncharacterized gene AGAP005440; in the other strain, the transgene was located on chromosome arm 2R at position 19768569 in an intergenic region (Figure 2B). These mosquito strains are referred to as AgY53B/2L and AgY53B/2R hereinafter.

In transgenic males, the activity of the CRISPR^AgY53B^ construct is hypothesised to shred the Y chromosome, leading to female bias in the progeny. To investigate this, we set-up a phenotypic assay experiment for both the mosquito strains, AgY53B/2L and AgY53B/2R. Males carrying the transgene in heterozygosity (AgY53B/2L +/- or AgY53B/2R +/-) as well as sibling males not harbouring the genetic construct (AgY53B/2L -/- or AgY53B/2R -/-) were mated *en masse* with WT females in independent cages. After blood feeding, females from each experimental sample were allowed to deposit eggs individually. For each female, the hatching rate (HR) was scored by counting the number of eggs laid and the larvae that hatched (Supplementary File S4). As showed in Figure 2C, strain AgY53B/2L +/- showed a significant reduction in HR when compared to the control AgY53B/2L -/- (HR median for AgY53B/2L +/- is 62.50 %, HR median for AgY53B/2L -/- is 80.82%). On the contrary, for strain AgY53B/2R, no significant differences in HR were observed when compared to the control (HR median for AgY53B/2R +/- is 88.30 %, HR median for AgY53B/2R -/- is 93.37 %). Progeny from each female was grown until the pupal stage, and the sex was assigned based on morphological observations. While progeny generated from the control strains showed a sex ratio around 50:50, progenies of AgY53B/2L +/- and AgY53B/2R +/- males showed a high level of female bias (median female bias = 98.75 % and 100 %, respectively) (Figure 2D).

Furthermore, when the fertility of female progeny was investigated, no fitness cost was detected, indicating that these females were fit and able to mate and give progeny (Supplementary Figure S1A and B and File S4). In addition, we set up a cross with AgY53B/2R +/- males and WT females to generate a large progeny and to be able to isolate any ‘escapee’ male harbouring or not the genetic construct (15 AgY53B/2R + and 15 AgY53B/2R - out of ∼3000 progeny analysed) (Supplementary Figure S1C and File S4). We performed a phenotypic assay on these males to investigate their fertility as well as the sex ratio in their progeny. We detected a slight, but significant decrease in the HR of progeny from AgY53B/2R + males (P value = 0,0448), while no reduction in HR was observed in the progeny from AgY53B/2R - males. Notably, AgY53B/2R + escapee males still produce highly female biassed progeny (Supplementary Figure S1D and File S4).

Taken together, these results indicated that targeting Y-linked satellite sequences using CRISPR/Cas9 does not impact the HR in the progeny of transgenic males, and it causes strong female bias in their progeny.

### 3 Shredding the Y chromosome during male gametogenesis causes anaphase chromosome lagging and sex chromosomes aneuploidy

Next, we investigated the cytogenetic mechanisms underlying sex ratio distortion due to Y chromosome shredding. To do this, we analysed meiosis in the testes of AgY53B/2L and AgY53B/2R males.

We performed DNA FISH using fluorescent probes designed to label the X-linked rDNA and the Y-linked satellite AgY53B. As shown in Figure 3A-B, sex chromosome specific probes allow to follow the behaviour of sex chromosomes during meiotic divisions in WT male mosquitoes. FISH experiments on Y-shredding strains showed no evident alterations in meiotic prophase and metaphase chromosomes (Supplementary Figure S2). However, as shown in Figures 3C and 3D, we identified the presence of DNA bridges/lagging in cells undergoing meiotic anaphase, which were not identified in WT controls. Y chromosome-specific signal (Figure 3C i-ii-iv and 3D i-ii-iv) localised with chromatin bridges, suggesting that anaphase defects are due to chromosome damage imposed by Cas9 endonuclease activity. Interestingly, throughout our observations, we noticed that, on rare occasions, the X chromosome signal was also found in chromatin bridges/lagging during meiotic anaphase (Figure 3C iii and Figure 3D iii). We extended FISH analyses to the *An. gambiae* strain, named Ag(PMB)1, that was previously developed to shred the X chromosome in male gametogenesis (Figure 5A shows the components of the genetic construct in this strain) (32). Our analyses revealed colocalization of the X-chromosome specific signal and DNA bridges/lagging, supporting the hypothesis that these anomalies are due to the damage caused by endonuclease activity (Supplementary Figure S3).

**Figure 3:**
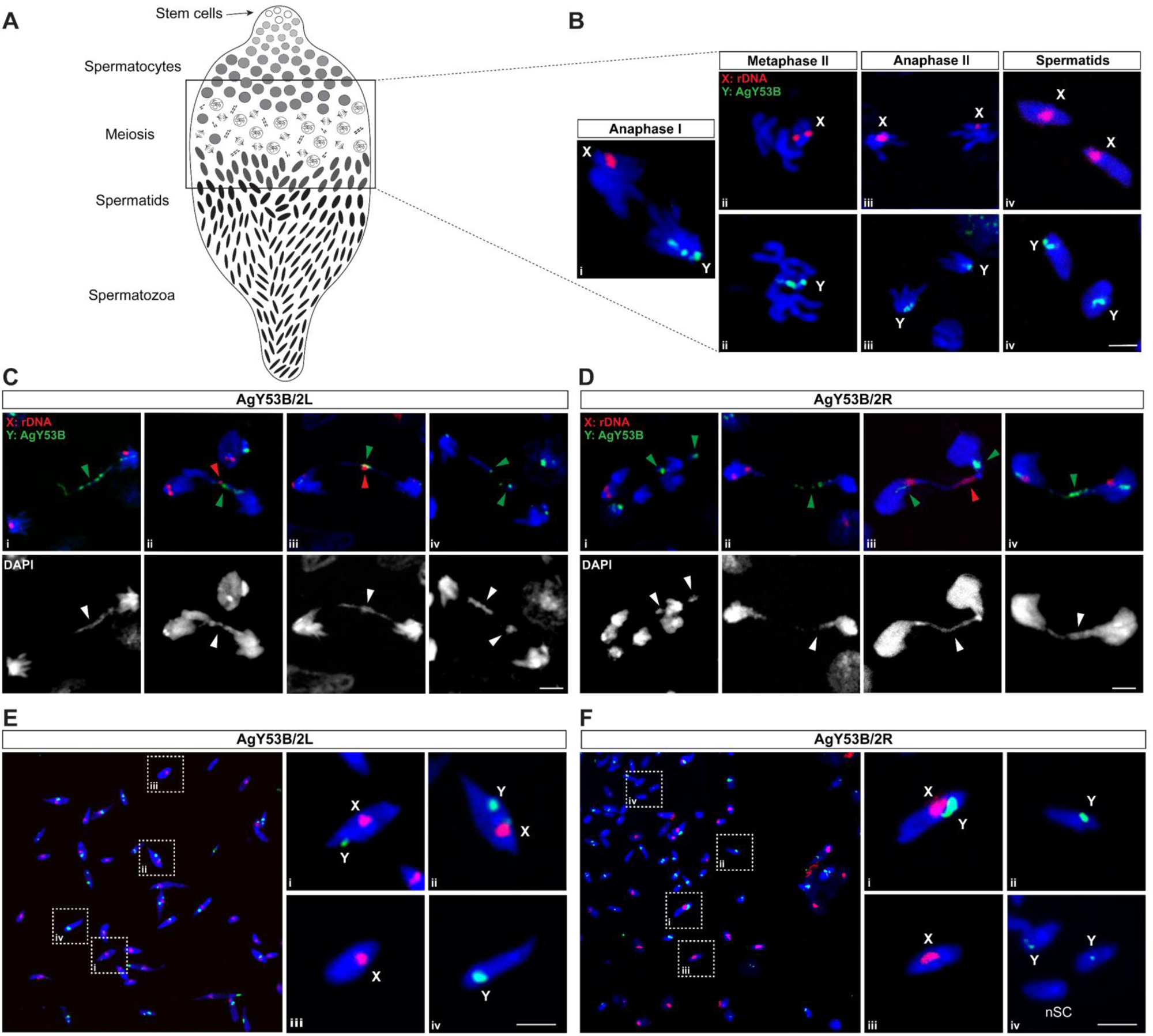
Cytological analyses of spermatogenesis in testes of Y chromosome shredding strains. (A) Schematic representation of the stages of spermatogenesis along the testis. (B) DNA FISH using probes specific to the X and Y chromosomes allows tracking of the sex chromosomes during meiotic stages in WT *An. gambiae*. Anaphase I is shown in panel B-I, where sex chromosomes segregate into two different cells. Panel B-ii shows meiotic metaphase II with chromosomes lined up along the metaphase plate. B-iii shows anaphase II where sister chromatids segregate. B-iv shows haploid spermatids resulting from the meiotic division. Panels C and E show meiotic defects observed in strain AgY53B/2L. Y chromosome lagging was observed in anaphase cells (C i-iv). In some cases, the X chromosome is also involved in lagging phenomena (C ii-iii). (E) Mature spermatozoa exhibit the signature of sex chromosome non-disjunction (NDJ). Mature spermatozoa with both X and Y chromosome-specific probes are shown in E-i and ii. Spermatozoa with the correct sex chromosome karyotype are shown in E-iii, iv. Similar meiotic defects were observed in strain AgY53B/2R (D-F). Y chromosome anaphase lagging is shown in D-i, ii, iv, X and Y chromosome lagging is shown in D-iii. F shows the sex chromosome karyotype in mature sperm. X-Y NDJ and sperm with no sex chromosome signal (nSC) are shown in panels i and iv. Sperm with normal sex chromosome karyotypes are shown in ii and iii. Red = X chromosome-specific probe (X: rDNA). Green = Y chromosome-specific probe (Y: AgY53B). Blue =DAPI. Scale bars = 3 µm. Green and red arrowheads indicate a lagging Y or X chromosome, respectively.

We then asked if the anaphase defects observed in the Y-shredding strains would result in miss segregation of the sex chromosomes in the following stages of meiosis. To answer this, we looked at Y-specific and X-specific signals in post-meiotic cells. In testis dissected from WT *An. gambiae*, the ratio of X and Y-bearing sperm is around 50 % (15,42). As shown in Figures 3E and 3F, both the Y-shredding strains analysed showed the simultaneous presence of both X and Y chromosome signals in mature sperm, suggesting the occurrence of X-Y nondisjunction events (X-Y NDJ). In addition, sperm lacking signals from the sex chromosome-specific probes (nSC) were detected, supporting the hypothesis of sex chromosome miss segregations during meiotic anaphases (Figure 3F iv). Sperm showing the expected only-X or only-Y chromosome segregation were also found (Figure 3E iii-iv, Figure 3F ii-iii). Surprisingly, a similar study on the Ag(PMB)1 strain showed that X-Y NDJ accounted for only 3.6 % of the total sperm counts (42).

Taken together, these results suggest that, despite HR being unaffected, shredding the Y chromosome during male gametogenesis leads to dramatic effects on sex chromosome segregation and gamete formation.

### 4 Sperm with sex chromosome non-disjunctions are negatively selected following Y chromosome shredding

We hypothesised that abnormal sperm would be removed at some stage of sperm production and/or prior to egg fertilization. This prompted us to further investigate the presence of abnormal sperm in the testis of Y shredding strains as well as in the spermathecae dissected from WT females after mating with transgenic males (Figure 4A).

**Figure 4:**
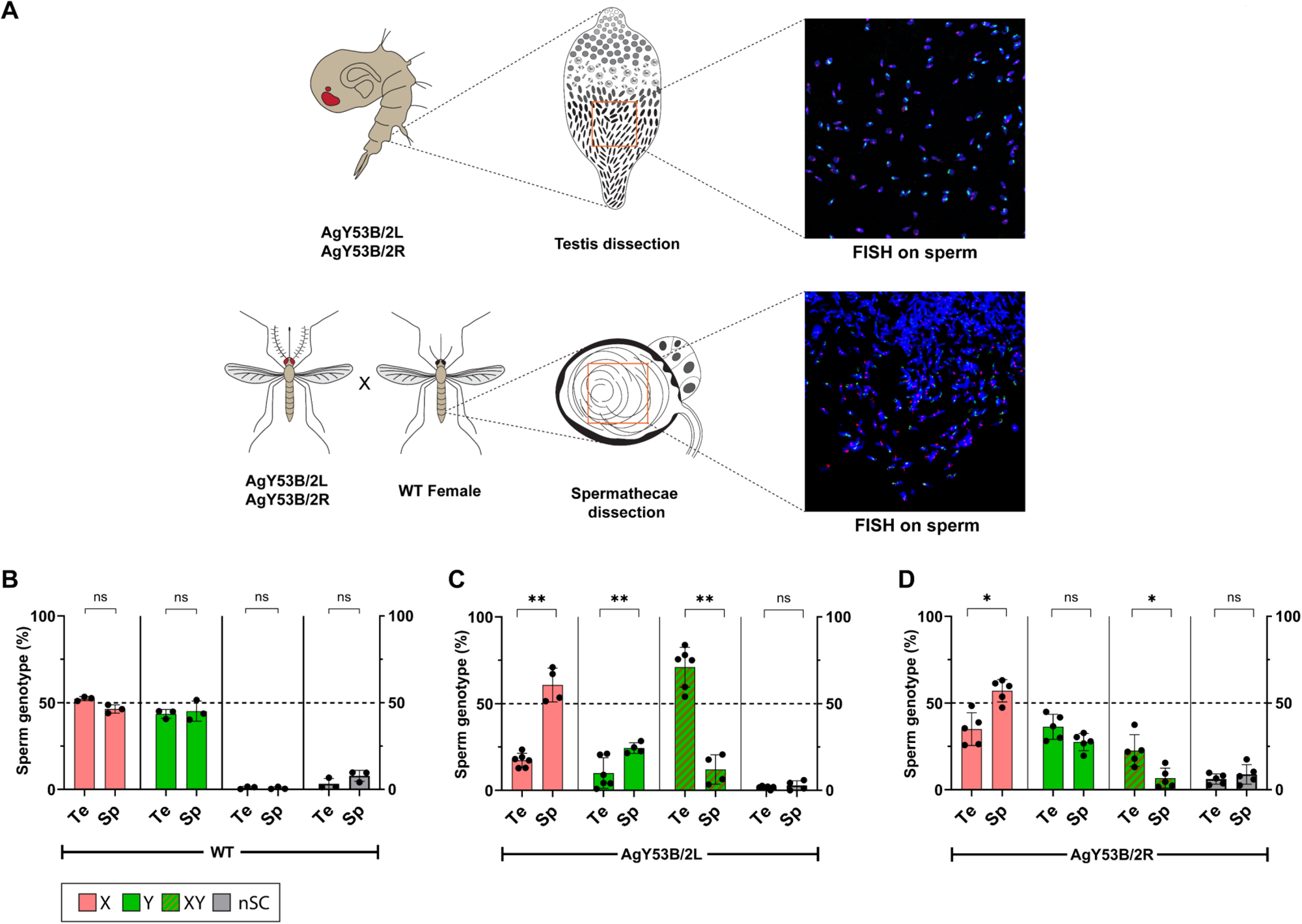
Investigating sperm selection in Y-shredding strains. (A) Schematic representation of the workflow used to perform sex chromosome counts in the sperm of WT and Y-shredding strains. FISH was performed with fluorescent probes targeting the X-linked rDNA locus (Red) and the Y-linked AgY53B satellite (Green). After confocal analyses, sperm counts were carried out using the Colocalization object counter in ImageJ (53). (B) The plot shows the ratio of sex chromosomes found in the testis (Te) of WT males and the spermathecae (Sp) of WT females mated with WT *An. gambiae* males. We classified the sperm based on 4 categories: X-bearing only (X), Y-bearing only (Y), X-Y non-disjunction (XY) and sperm with no sex chromosome signal (nSC). No significant difference in the ratio of sex chromosomes was detected in both the testis and the spermathecae. Number of Testis (Te) = 3, Number of Spermathecae (Sp) = 3. Testis total sperm counts: X = 760, Y = 626, XY = 14, nSC = 57. Spermathecae total sperm counts: X= 371, Y = 365, XY = 5, nSC = 63. (C) Ratio of sex chromosomes found in the testis of AgY53B/2L males. Testes of these males show a high proportion of X-Y NDJ compared to X or Y only bearing sperm. Sperm counts on female spermatheca inseminated by AgY53B/2L males show a significant reduction in the ratio of X-Y bearing sperm, while X and Y only bearing sperm show a significant increase in their ratio (Mann Whitney test, *P value* < 0,01 for all conditions). We found no significant change in the ratio of nSC sperm between the counts performed on the testis and spermathecae. n Te = 6, n Sp = 4. Te total sperm counts: X = 145, Y = 89, XY = 561, nSC = 14. Sp total sperm counts: X = 414, Y = 177, XY =68, nSC = 19. (D) Ratio of sex chromosomes found in the strain AgY53B/2R. This strain exhibits a lower number of X-Y NDJ in the testis compared to AgY53B/2L (median 21.88 % vs 75.27 %). The ratio of sperm showing X-Y NDJ significantly decreases following mating and storage in females’ spermathecae (Mann Whitney test, *P value* < 0,05). We observed a statistically significant increase in the ratio of X-bearing sperm in the spermatheca compared to the testis (Mann Whitney test, *P value* < 0,05). We found no significant differences in the ratio of Y-bearing and nSC sperm between the testis and spermathecae in this strain. n Te = 5, n Sp = 5. Testis total sperm counts: X= 782, Y = 896, XY = 474, nSC = 180. Spermathecae total sperm count: X = 1005, Y = 592, XY = 69, nSC = 94. In each bar, the median with Standard Deviation (SD) is shown.

For this study, we relied on DNA FISH and sex-chromosome-specific probes, which allowed us to conduct a quantitative analysis of normal *versus* abnormal sperm in WT and experimental samples. In the reproductive organs dissected from WT pupae, a ∼50 % ratio of only-X and only-Y-bearing sperm was observed over a sample size of 1457 sperm collected from 3 different individuals. Similar results were found in the spermathecae dissected from WT females mated to WT males (sample size = 804 from 3 spermathecae). In WT males, abnormal sperm (X-Y NDJ and nSC) were detected at a frequency of 1.5 % in the testis, while X-Y NDJ and nSC sperm were detected at 0.37 % and 9.5 % in the spermathecae of WT females, respectively (Figure 4B). Sex chromosome counts in the testes of Y-shredding strains and the spermathecae of females inseminated by these males showed an enrichment of only-X-bearing sperm in the latter, with a bigger increase in strain AgY53B/2L (Figures 4C and 4D). The proportion of only-Y bearing sperm increased in the spermatheca compared to the testis, when females were mated with AgY53B/2L males, while a non-significant difference was observed for strain AgY53B/2R (Figure 4C-D). Interestingly, although a high proportion of XY-bearing sperm was observed in the testis of both Y-shredding strains (median X-Y NDJ AgY53B/2L = 75.27 %, AgY53B/2R = 21.88 %), this number dramatically decreased in the spermathecae of females inseminated by these males (median X-Y NDJ AgY53B/2L = 12.9 %, AgY53B/2R = 4.54 %) (Figure 4C-D). Counts from testes and spermathecae and statistical tests can be found in Supplementary File S6.

Taken together, these results suggest a strong negative selection on sperm, showing X-Y NDJ signatures that take place between sperm production and sperm storage in the female spermathecae. On the contrary, a positive selection acts on only-X or Y-bearing sperm, with a higher proportion of only-X-bearing sperm found in spermathecae; this is in line with the high HR observed for the Y-shredding strains and the high level of female bias detected in their progeny. However, a relatively high proportion of Y-bearing sperm is still present in the spermathecae of these females after insemination, suggesting the presence of a different mechanism of sperm selection that will require further characterization.

### 5 Investigating the effects of the simultaneous shredding of X and Y chromosomes during male gametogenesis in *An. gambiae*

The *An. gambiae* sex chromosomes exhibit significant differences in terms of size, genetic composition, and gene content. We hypothesised that the simultaneous shredding of the X and Y chromosomes in male gametogenesis might reveal distinct chromosome behaviour in response to damage. This could have an impact on meiotic divisions, sperm selection, male fertility, and potentially the sex ratio in the progeny.

To investigate this, we set up genetic crosses between the Y-shredder strains AgY53B/2R or AgY53B/2L, and the previously established X-shredder strain Ag(PMB)1 (32,54). We generated trans heterozygous males harbouring the genetic construct targeting the X chromosome (inherited from Ag(PMB)1 parental males) in combination with the genetic construct targeting the Y chromosome (inherited from either AgY53B/2L or AgY53B/2R parental females) (Figure 5A). Subsequently, we conducted a phenotypic assay by crossing WT *An. gambiae* females with Ag(PMB)1-AgY53B/2L or Ag(PMB)1-AgY53B/2R males. As shown in Figure 5B, for Ag(PMB)1-AgY53B/2L and Ag(PMB)1-AgY53B/2R we observed a strong reduction in HR. Contrary to this, no reduction in HR was detected in the parental Y shredding strains (Figure 2C), and in the Ag(PMB)1 strain, used as a control in this experiment (HR median for Ag(PMB)1-AgY53B/2L is 34.78 %, HR median for Ag(PMB)1-AgY53B/2R is 37.50 % and HR median for Ag(PMB)1 is 90 %). Surprisingly, when we scored the sex ratio in the progeny of the trans heterozygous males, a strong female bias (94.28 % and 85.71 %) was observed, while for the Ag(PMB)1 strain, a high male bias was observed in line with previous data (Figure 5B, (32)).

**Figure 5:**
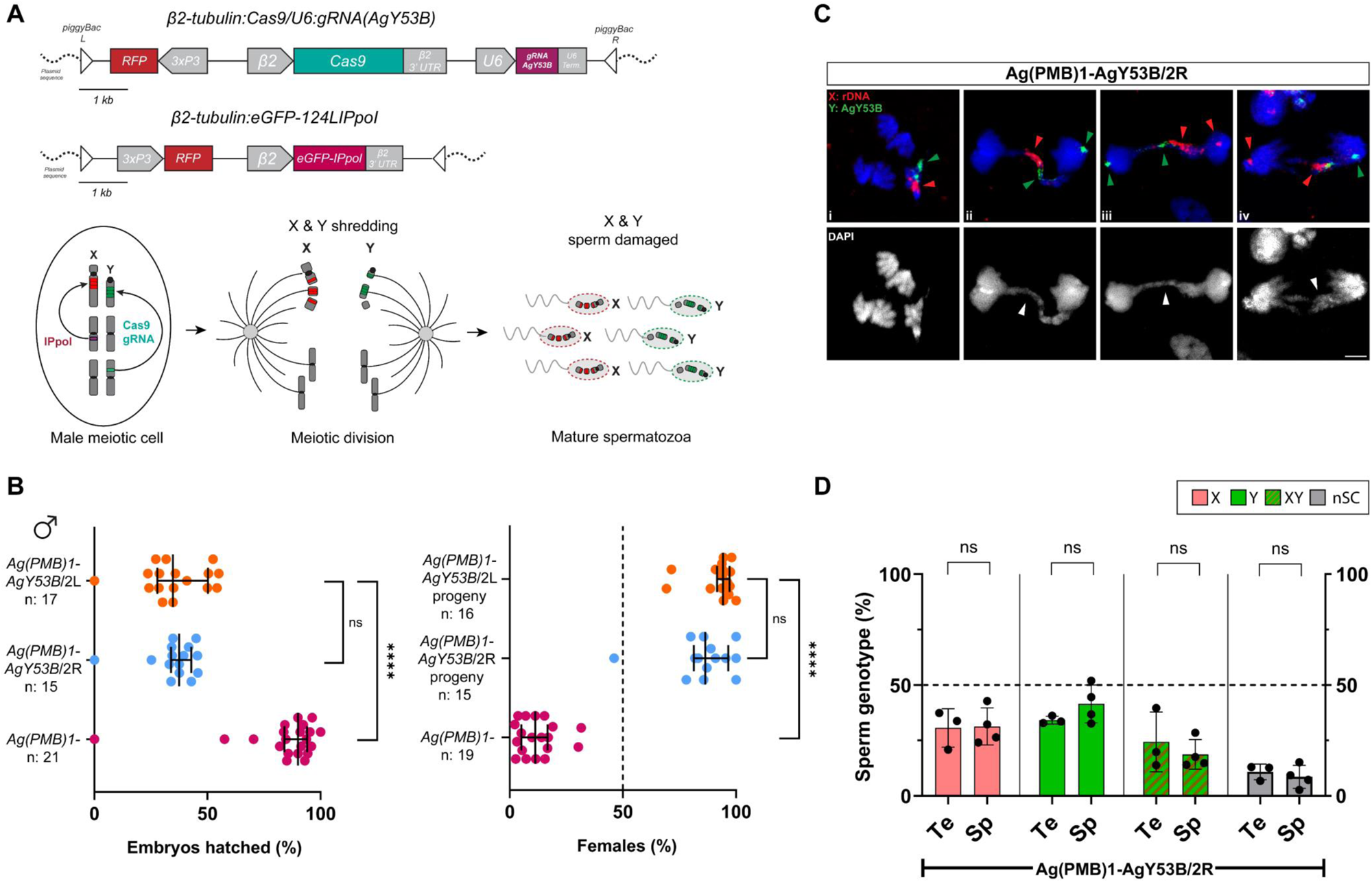
Investigating the effects of simultaneous X and Y chromosomes shredding on HR, sex ratio, meiosis and sperm selection. (A) Graphic representation of the genetic constructs used to induce Y (top) and X (bottom) chromosome shredding and expected nuclease activity during male meiosis. (B) Effects of simultaneous X and Y chromosomes shredding. Note that in this experiment, strain Ag(PMB)1 is used as a control. Simultaneous X and Y chromosome shredding causes a strong reduction in HR (HR median from top to bottom, 34.78 %, 37.50 %, 90 %). No significant difference in the reduction of the HR was detected between the two trans-heterozygous strains tested (Dunn’s multiple comparison, *P value* > 0,999). We found a high level of female bias in the progeny of both the X-Y shredding strains. No statistical difference was detected in the level of female bias between the two trans-heterozygous Ag(PMB)1-AgY53B strains (non-parametric t-test, *P value* > 0,05). The median of the percentage of females from the progenies is as follows: 94.28 %, 86.41 % and 11.25 % (from top to bottom). (C) DNA FISH on strain Ag(PMB)1-AgY53B/2R reveal lagging phenomena involving both sex chromosomes. In panels C-ii, iii, iv red and green arrowheads indicate partially segregating and lagging sex chromosomes. Panel C-i shows an early stage of anaphase where autosomes start to segregate and sex chromosomes are still paired. Grey arrowheads indicate chromatin bridges/lagging chromosomes. Red = X chromosome-specific probe (X: rDNA). Green = Y chromosome-specific probe (Y: AgY53B). Scale bars = 3 µm. (D) Investigation of sperm selection in strain Ag(PMB)1-AgY53B/2R. We observed a similar ratio of X and Y bearing sperm in the testis dissected from Ag(PMB)1-AgY53B/2R (median of 33 %). In the testis, the ratio of sperm exhibiting X-Y NDJ has a median of 19.75 % while sperm with nSC were 12.65 %. We observed no significant differences in the ratio of sperm sex chromosome genotype between testis and spermathecae (Mann Whitney test, *P value* > 0,05). N Te = 3, n Sp = 4. Testis total sperm counts: X = 327, Y = 361, XY = 254, nSC = 116. Spermathecae total sperm count: X = 467, Y = 537, XY = 252, nSC = 119. In each bar, median with Standard Deviation (SD) is shown.

These results suggest that despite simultaneous shredding of the sex chromosomes, functional sperm are produced that are capable of fertilising eggs upon mating. In addition, X-bearing sperm are favoured over Y-bearing sperm.

Next, we performed DNA FISH on testis dissected from trans heterozygous Ag(PMB)1-AgY53B/2R males to investigate the behaviour of the sex chromosomes during meiosis following X-Y shredding. As shown in Figure 5C, we observed DNA bridges/lagging of both sex chromosomes in meiotic anaphase I. The sex chromosome lagging pattern in anaphases shows that a large portion of the sex chromosomes lags between the two dividing cells. However, a smaller portion of the sex chromosomes partially segregates at the poles of the meiotic spindle (Figure 5C ii-iii-iv). The same pattern of anaphase lagging was observed in testis dissected from trans heterozygous Ag(PMB)1-AgY53B/2L males (Supplementary Figure S4).

The investigation of sperm selection through DNA FISH was carried out only in trans heterozygous Ag(PMB)1-AgY53B/2R males. We quantify the sex chromosome karyotype in testes dissected from these males and the spermathecae of females after mating, as previously described. Surprisingly, we found no significant difference in the ratio of sex chromosomes in sperm from testes and female spermathecae (Figure 5D). In addition, we found that female spermathecae had a similar proportion of X- and Y-only bearing sperm (median of the percentage, X = 29.26 % and Y = 40.64 %). Interestingly, the number of X-Y NDJs was higher when compared to the spermathecae of females mated with Y shredding males (median of the percentage X-Y NDJ Ag(PMB)1-AgY53B/2R = 16.13 % versus AgY53B/2R = 4.54 %) (Figure 4-D and Figure 5D). A small proportion of sperm lacking signal from sex chromosome-specific probes was also observed in both testes and spermathecae (median of percentage nSC, Tesits = 12.6 % versus Spermathecae = 8.2 %).

Taken together, these results suggest that targeting both X and Y chromosomes during male gametogenesis does not result in full sterility but significantly impacts male fertility. Our analyses reveal that, although meiotic defects are present, functional sperm are produced. Lastly, our findings show a high level of sex ratio distortion towards females in the progeny of trans heterozygous males, indicating a competitive advantage of X-bearing sperm over Y-bearing sperm. Additional experiments to score sperm sex chromosome genotypes in trans heterozygous Ag(PMB)1-AgY53B/2L testis are provided in supplementary material (Supplementary Figure S4-B).

### 6 Shredding the Y chromosome at early stages of embryogenesis and spermatogenesis causes male-embryo lethality and Y chromosome loss, respectively

The generation of a Y-shredder system allows for a functional study of the Y chromosome in a range of tissues and organs and at different stages of development. We made use of the *Vasa2* promoter, active in the germline of both sexes and proven to induce maternal Cas9 deposition (52), to investigate the outcome of early embryo Y chromosome shredding (52).

We generated trans heterozygous females harbouring *Vasa2*:Cas9 in combination with CRISPR^AgY53B^ (Vasa2:Cas9-AgY53B/2R) and mated them with WT males (Figure 6A). After mating and blood feeding, females were allowed to lay singularly, and their HR was measured and compared to control sibling females only harbouring the CRISPR^AgY53B^ construct. In the progeny of *Vasa2*:Cas9-AgY53B/2R females, we observed a median HR of 43.05 % with almost only females recovered at the adult stage (336 out of 339). Conversely, an HR of 89.79 % was found in control females with the expected 50:50 sex ratio in the progeny (Figure 6B, Supplementary File S5). A reduction in the HR and a high level of female bias in the progeny of *Vasa2*:Cas9-AgY53B/2R females demonstrate that maternal deposition of Cas9 and gRNA targeting satellite AgY53B leads to male-specific embryo lethality.

**Figure 6:**
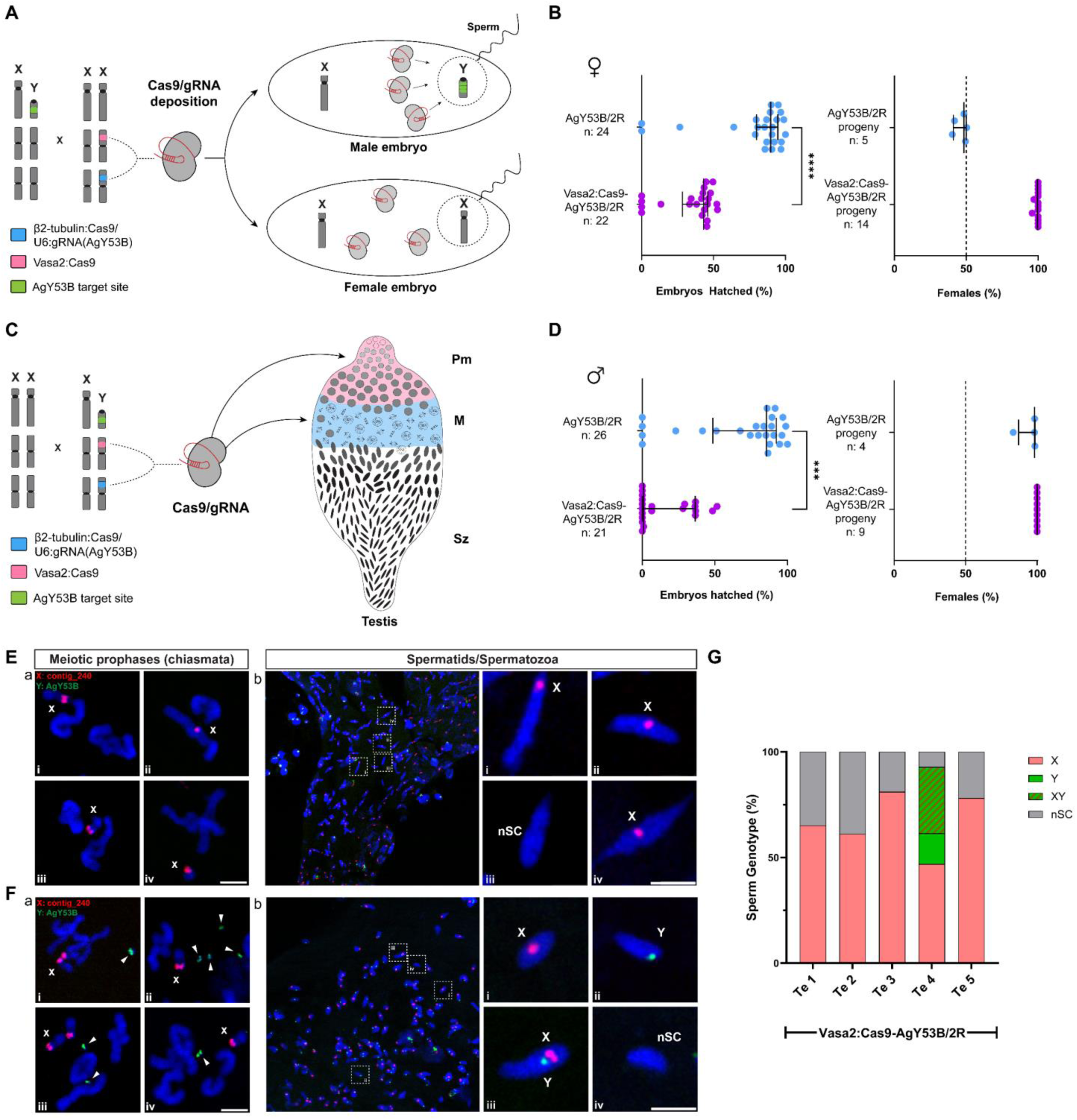
Y chromosome shredding in early embryos and at early stage of spermatogenesis. (A) Graphic schematic showing how Cas9 expression under Vasa2 promoter in parental females is expected to cause deposition of Cas9 and male-embryo lethality. (B) Maternal deposition of Cas9/gRNA targeting the Y chromosome causes a nearly 50 % reduction in HR (median HR from top to bottom: 43.05 %, 89.79 %, non-parametric *t-test*, *P value* < 0,05) and strong female bias in the progeny. Sibling females used as controls (AgY53B/2R) have no reduction in HR and show the expected sex ratio of 50 %. (C) A graphic schematic showing the expression pattern of Cas9 in trans heterozygous males (Vasa2:Cas9-AgY53B/2R). Pre-meiotic cells = PM, Meiotic = M, Spermatozoa = Sz. (D) Trans-heterozygous Vasa2:Cas9-AgY53B/2R males show a strong reduction in HR following the activity of Cas9 expressed under Vasa2 promoter when compared to the control males harbouring only the CRISPR^AgY53B^ construct (median HR from top to bottom, control: 85.55 %, trans heterozygous males: 0.96 %, non-parametric *t-test*, *P value* < 0,05). Control males produce a high female biassed progeny (median 98.20 %) due to the activity of CRISPR^AgY53B^ construct. No males were detected in the progeny of Vasa2:Cas9-AgY53B/2R males. (E) DNA FISH on testes from Vasa2:Cas9-AgY53B/2R males reveal loss of the Y chromosome during meiosis and absence of Y-bearing sperm. E-a panels show meiotic prophase chromosomes (presence of chiasmata), from which only the X chromosome is detectable. E-b panels show the sex chromosome karyotype of sperm based on the signal retrieved from sex chromosome specific probes. Only X-bearing or sperm with no sex chromosomes are detectable as a product of spermatogenesis. The presence of a green signal (Y-probe) in panel E-b is likely to derive from either somatic cells surrounding the testis or the Male Accessory Glands (MAGs). (F) DNA FISH on testes from Vasa2:Cas9-AgY53B/2R males reveal fragmentation of the Y chromosome and the presence of Y bearing sperm. F-a panels show meiotic prophase chromosomes, from which it is possible to detect signals from the Y chromosome specific probe (green arrowheads). The Y chromosome appears fragmented and smaller in size when compared to AgY53B/2L and AgY53B/2R strains (Supplementary Figure S6). F-b panels show the sex chromosome karyotype in the sperm of trans heterozygous males. In this case, sperm show a signal from the Y-chromosome specific probe (panel F-b-ii). X-bearing, X-Y NDJ and nSC sperm are also present (F-b-i-iii-iv). Chromosomes and sperm shown in panels F-a and F-b were obtained from the same testis. (G) Sex chromosome genotyping of sperm from 5 Vasa2:Cas9-AgY53B/2R males. 4 out of 5 males show only X-bearing or sperm bearing no sex chromosomes. In all these males, X-bearing sperm account for more than 50 % of the total sperm count. One testis (Te-4) shows the presence of X, Y, XY and sperm with no sex chromosomes (nSC). Counts performed on Te-4 were performed on the same testis shown in F-a and F-b. Total sperm counted (n) from each individual testis (Te) are, from left to right, 201, 126, 217, 394, 55. Blue = DAPI. Red = X chromosome-specific probes (X: conting_240). Green = Y chromosome-specific probe (Y: AgY53B). Scale bars = 3 µm.

Next, we generated trans heterozygous males harbouring *Vasa2*:Cas9 in combination with CRISPR^AgY53B^. This genetic background allows us to evaluate the impact of Y chromosome shredding at an early stage of spermatogenesis (pre-meiotic cells) (Figure 6C). We set up crosses between *Vasa2*:Cas9-AgY53B/2R males and WT females. After mating and blood feeding, females were allowed to lay singularly, and their HR was measured and compared to control sibling females mated with males only harbouring the CRISPR^AgY53B^ construct. We observed a strong reduction in the HR (median 0.96 % and mean 15 %) in the progeny of females mated with trans heterozygous males, with only females recovered at the adult stage (Figure 6B, supplementary File S5). Conversely, control females mated with AgY53B/2R males showed a median HR of 85.55 % and a strong female bias (median 98.2 %) in the progeny (Figure 6-D).

The recovery of progeny from males that undergo pre-meiotic shredding of the Y chromosome in their testes prompted us to investigate this phenotype more in detail. First, we dissected the spermathecae of females mated with trans heterozygous *Vasa2*:Cas9-AgY53B/2R males. Out of the 25 spermathecae recovered from females that laid eggs, 21 contained sperm, suggesting that the mating ability of trans heterozygous males is not impaired (Supplementary File S5).

Next, we wanted to investigate the effect of pre-meiotic Y-chromosome shredding on the reproductive organs of these males at both macro- and microscopic level. We performed DAPI staining on testes dissected from five trans heterozygous males. We observed a range of phenotypes among the testes dissected. Some testes were atrophic, small, and had only a few cells populating the organ (Supplementary Figure S5). Conversely, some testes showed a normal-like shape, where all the spermatogenesis stages were detectable and mature spermatozoa were visible, however, as shown in Supplementary Figure S5, DAPI staining revealed that normal-like testis had very large spermatid nuclei with an elongated shape. In addition, mature spermatozoa had an aberrantly curved shape when compared to the arrow-like shaped sperm found in the testis dissected from WT control or from AgY53B/2R strain (Supplementary Figure S5). Furthermore, we observed a lower number of mature sperm in testes with normal shape compared to the controls (Supplementary Figure S5, WT and AgY53B/2R, testes dissected from one day old adult males).

To investigate sex chromosome behaviour in trans heterozygous *Vasa2*:Cas9-AgY53B/2R males, we performed DNA FISH using sex chromosome-specific probes as previously described. Following confocal analyses, we found two different scenarios that we categorised as Y-loss (Y-L) and Y-partial-loss (Y-P-L). In the Y-L category, shown in Figures 6E-a-b, we observed only X-specific probe signal from prophase meiotic chromosomes (Figure 6E-a i-ii-iii-iv). We did not observe mature sperm with Y-specific probe signal in testes dissected from this category (n = 4) (Figure 6E-b i-ii-iii-iv and Figure G). Contrary, in the Y-P-L category, the presence of the Y chromosome signal was observed (Figure 6F-a i-ii-iii-iv). However, the signal revealed Y chromosome fragmentation and the absence of X-Y chromosome pairing. In addition, the Y chromosome is smaller in size when compared to AgY53B/2R or AgY53B/2L strains (Supplementary Figure S6).

To estimate the occurrence of the sex chromosome categories identified, we scored sex chromosome genotypes in sperm from 5 trans heterozygous *Vasa2*:Cas9-AgY53B/2R one day-old adult males (Supplementary File S6). As shown in Figure 6G, 4 out of 5 of the testes did not show Y-bearing sperm but only X-bearing or sperm with no sex chromosomes (nSC). In one male, we found the presence of Y bearing sperm (counts were performed on the same testis shown in Figure 6F) in concomitance with X-Y NDJ sperm and nSC sperm.

Taken together, these results suggest that although pre-meiotic shredding of the Y chromosome dramatically affect fertility of males, mature sperm can still be produced. As a result of the pre-meiotic shredding, partial or entire loss of the Y chromosome can be observed in these spermatozoa. The presence of testes where only X-bearing sperm were detected suggests that meiosis can proceed despite the loss of the male chromosome, and these sperm are able to fertilise eggs, leading to only female progeny. The absence of male progeny suggests that even if Y-bearing sperm are produced in some cases, they are not able to produce viable individuals.

## Discussion

Through our analyses, we identified Y-chromosome-specific DNA sequences suitable for the development of a CRISPR/Cas9-based SSD system in *An. gambiae*. We selected two target sites, gRNA-AgY53A and gRNA-AgY53B, and validated the Y-linkage through DNA FISH experiments, providing insight into the organisation of such elements along the chromosome length. A similar approach could be used to virtually map any Y-linked sequence identified through this study, with the potential to generate a high-resolution cytogenetic map of the Y chromosome in *An. gambiae.* This could be valuable to investigate the spatial distribution of these genetic elements and their role in biological mechanisms, such as sex chromosome pairing.

We used a CRISPR/Cas9 system to target the satellite AgY53B, and we generated two sex distorted strains, AgY53B/2L and AgY53B/2R, with the latter showing a low reduction in fitness and the former showing no apparent fitness costs. This difference could be attributed to positional effects influencing the level of expression of Cas9. Alternatively, in strain AgY53B/2L, intragenic insertion of the genetic construct could affect the activity of the endogenous gene AGAP005440, which could be involved in mitotic division (55).

Both strains showed a strong female bias in the progeny, with both females (> 95 %) and males (< 5 %) able to mate and produce progeny. Interestingly, the Y chromosome of escapee males can undergo subsequent rounds of shredding, in line with observations previously made on escapee females from the X shredding strain Ag(PMB)1 (32).

The level of sex distortion observed in this study suggests that a similar approach could be used to develop Y shredding systems in other organisms even when the assembly of the Y chromosome is not complete. This would be of interest for species, such as vertebrate pests, where introducing a strong female bias has been theoretically proven to be a valuable strategy for population control (43,56).

The cytological investigation of the reproductive organs in the Y shredding strains highlighted defects in chromosome segregation identifiable as DNA bridges/lagging in meiotic anaphases. These abnormalities colocalised with the fluorescent probe designed to label the Y-linked target site, indicating an association with Cas9 activity. This is in line with previous studies in *Drosophila* showing that X chromosome lagging can be induced by the activity of I-Crel endonuclease in somatic tissue (57).

Interestingly, in some cases, the signal used to label the X chromosome colocalised with the lagging structures, suggesting an interaction between this chromosome and the Y-linked target site. This is supported by a study showing that satellite elements, named ‘AgY477-AgY53B junction’, and containing partial sequences of satellite AgY477 and AgY53B, can be found in the centromeric region of the X chromosome in species of the *Anopheles gambiae* complex (26). Satellite elements are known to be the main components of centromeric DNA and play a key role in ensuring correct chromosome segregation (58–60). In addition, traces of recombination between the sex chromosomes in *An. gambiae* have been found to occur between genomic regions harbouring satellite elements (14).

In line with the abnormalities observed, when we investigated post-meiotic sperm cells, non-disjunction events of sex chromosomes were identified. Similar chromosome segregation defects have also been observed in natural sex ratio meiotic drives, especially among *Drosophila* and mosquito species (61–66).

The cytological defects that we observed in the Y shredder strains do not reflect on the overall high HR we recorded. This can be explained by the presence of mechanisms for sperm selection that we have shown take place prior to fertilisation. Although the molecular pathways regulating these selection mechanisms were not investigated, we hypothesise that the aberrant amount of DNA in sperm harbouring NDJ events could interfere with histone-to-protamine transition during spermiogenesis, impairing full sperm maturation. Recently, sperm elimination in the presence of a segregation distorter meiotic drive has been investigated in *D. melanogaster.* In this scenario, the severity of spermatogenic defects is linked to the copy number of the meiotic drive target, the *Responder* (Rsp) satellite, and in the presence of a high copy number of *Rsp*, checkpoints are activated during the histone-to-protamine transition, leading to cell death (64). TUNEL assays to investigate if the apoptotic pathway is activated in the Y shredder strains could be conducted in the future (67).

Despite the presence of negative selection, our analyses reveal that a low percentage of XY NDJ sperm can still be transferred to the female spermathecae upon mating. Female *Anopheles* mosquitoes mate mostly once in their lifetime and store the sperm in the spermatheca. Sperm are then used to fertilise the eggs during oviposition, following a blood meal. Is it likely that when females are inseminated by males that produce defective sperm, viable sperm are the first to be used for fertilisation, and the presence of a high level of HR in our Y shredding system points towards this hypothesis. A possible experiment to test if defective sperm can eventually be used by females to fertilise their eggs would be forcing the use of sperm until storage exhaustion, which could be achieved through multiple gonochoric cycles.

Furthermore, despite the high female bias in the progeny of Y shredding males, Y-bearing sperm can still be found in the female spermathecae upon mating, and it would be interesting to explore if, through multiple gonochoric cycles, their appearance in the progeny would increase.

We then investigated the consequences of simultaneous shredding of the sex chromosomes, which strongly reduced the ability of males to produce viable sperm and, interestingly, led to a strong female bias in the progeny. We hypothesised, in this scenario, that a strong positive selection acts on the only-X and only-Y-bearing sperm that escape the shredding (estimated to be: ∼1% for the Y-shredding system and ∼5% for the X-shredding system), with an advantage for X-bearing sperm over Y-bearing sperm. This might be due to differences in the molecular mechanisms that are triggered upon X or Y chromosome shredding. For example, studies in *Drosophila* and yeast revealed that the rDNA locus is highly dynamic and undergoes a phenomenon called ‘rDNA magnification’ that causes the rapid expansion of rDNA copy number in the male germline in individuals with large rDNA deletions (68–71). This process is achieved by a mechanism named unequal sister chromatid recombination/exchange, where new copies of rDNA can be created by homologous recombination between sister chromatids following double-stranded breaks (DBSs) at the rDNA locus (70). Recent studies in *Drosophila* show that this dynamic can also occur in the male germline through ageing (71). We can hypothesise that the mechanism of copy number recovery could be responsible for the higher tolerance of Cas9-mediated DSBs at the rDNA locus, which could contribute to increased fitness of the X chromosome-bearing sperm when compared to the Y-bearing sperm and, ultimately, be responsible for the female bias observed in the progeny.

Furthermore, we demonstrated that maternal deposition of Cas9 in the presence of the gRNA targeting the Y chromosome can selectively kill male embryos. Importantly, this result supports Y chromosome specificity for the target sites selected (no off-targets). It is known that the male determining factor, *Yob,* is expressed 2 hours after egg-laying, while expression mediated by the *Vasa* promoter is no longer detectable after 2 hours of development (21,52). The death of male embryos could be explained by early shredding of the Y-chromosome (within the first 2 hours of development) interfering with the expression of *Yob* and potentially impairing activation of the dosage compensation pathway, in line with what was observed in Krzywinska et al. 2016 (21); notably, other Y-linked genetic elements might play a crucial role at this stage of embryonic development. Alternatively, male embryonic death could simply be the response to chromosomal damage.

We have also generated trans-heterozygous males expressing Vasa2-Cas9 in the presence of the Y shredding construct. *Vasa2* promoter drives Cas9 expression in pre-meiotic stages of gametogenesis, and somatic leakiness for this promoter has been previously reported (72–74). A recent experiment conducted in our laboratory (Silvia Grilli, personal communication) showed that trans-heterozygous individuals expressing Vasa2-Cas9 in the presence of the X shredding construct die at L3/L4 larval stage, suggesting that leaky expression of Cas9 can lead to mortality when targeting an essential locus. Notably, this is in contrast with what was reported in Galizi et. al. 2016 (33) where lethality was not reported as observations were made at an earlier developmental stage. The viability of trans-heterozygous males expressing Vasa2-Cas9 in the presence of the Y shredding system suggests that targeting the Y chromosome using a pre-meiotic and leaky promoter does not cause a toxic effect. This could be due to the scarcity of genes present on the Y chromosome that might not have a prominent role in the somatic tissues where Vasa2 promoter is leaky. However, although we could retrieve viable trans-heterozygous individuals that developed to the adult stage, we observed a strong reduction in their fertility (HR 0.96 %). This was reflected by the presence of atrophic testes in the experimental males, which highlights that pre-meiotic Y chromosome shredding can have a profound impact on male germline development. However, in some males, normal-like testes were retrieved, and meiotic chromosomes and mature sperm were observed with a range of chromatin condensation defects. This variable phenotype could be attributed to individual-specific variations in the level of expression and leakiness of Vasa2 promoter, which can result in a range of germline cellular defects. The cytogenetic analyses performed on normal-like testis revealed that pre-meiotic shredding can cause partial or entire loss of Y chromosome during spermatogenesis without impairing the progression of meiosis, leading to the production of viable X-bearing sperm, albeit at a notably low rate, which can still fertilise the eggs. This is in contrast with recent studies in *Drosophila* showing that Y-linked genes are essential for male fertility, as their knockout causes complete male sterility (75,76). Our experiment suggests that the *An. gambiae* Y chromosome might be dispensable for male gametogenesis. However, further experiments to accurately investigate the role of Y-linked elements in the reproductive tissues would be required.

In this study, we have developed the first Y shredding system in *An. gambiae*. This mosquito strain could be used as a genetic sexing system to produce a female biased progeny, which could support the development of research projects based on this organism. For instance, the features of this strain, in combination with the X-shredder strain, could allow high-throughput sex-specific transcriptome analyses of early embryos. A similar strategy, based on the use of natural occurring sex ratio meiotic drives, has recently been employed in *Drosophila* to explore the sex determination pathway (77,78) which, in *An. gambiae*, has only been partially characterised (79). Genes involved in sex determination are crucial for the development of genetic strategies aiming at vector control, thus meriting interest (31,35). Our study provides proof of principle that Y chromosome shredding can be tolerated in specific tissues and at specific developmental stages without causing lethality. These findings encourage further investigation of the effects of Y chromosome loss in tissues where it might play a significant role, such as those involved in mating behaviour (80). Altogether, this research enhances our general understanding of Y chromosome biology, and it could represent valuable knowledge for the development of novel control strategies aimed, for example, at impairing mating behaviour (81).

## Materials and Methods

### 1 Mosquito strains and maintenance

In this work, transgenic *An. gambiae G3* strains, previously generated in our lab, were used. Strain named Ag(PMB)1 harbours a genetic construct where beta 2 tubulin promoter mediates the expression of the I-Ppol structural variant W124L fused to eGFP during male spermatogenesis. Pax (3xP3) promoter drives the expression of RFP marker allowing the identification of transgenic individuals. The expression of I-Ppol endonuclease in male germline leads to approximately 95 % male offspring (32). The transgene is located on Chromosome 2R band 19D (82). The strain named Vasa2:Cas9 harbours a genetic construct where Vasa2 5’ regulatory sequences mediate the expression of Cas9 endonuclease in the germline of both male and female mosquitoes (52,83). Yellow Fluorescent Protein (YFP) marker is driven by Pax (3xP3) promoter, allowing for transgenic identification. The Vasa2:Cas9 genetic construct is integrated into chromosome 2R, the exact genomic coordinates are unknown. *An. gambiae* G3 was used to generated both AgY53B/2L and AgY53B/2L strains. All mosquitoes used in this study were maintained under standard mosquito-rearing conditions at 27°C ± 1 and 70 % ± 5 % relative humidity.

### 2 In *Silico* identification of Y-specific target sites

A modified version of Redkmer pipeline (46) was used to discover Y-specific gRNAs target sequences. Long-read whole genome sequencing (WGS) data were generated using Oxford Nanopore Technology (ONT) from genomic DNA extracted from 5 males and 5 females *An. gambiae* G3 strain. Genomic extractions, sequencing using MinION platform and quality checks were performed according to Vitale et. al. 2022 (82). Only Nanopore reads between 1kbs and 100kbs were selected as input for RedKmer pipeline. Illumina male and female WGS short reads were obtained from SRA archive under the project name PRJNA397539. Short-read raw data quality checks were performed according to Vitale et. al. 2022 (82). 5 male and 5 female-specific short-read data derived from a single adult mosquito were merged to obtain a minimum coverage of 20X. Long-read (pooled males and females) ONT sequencing data are available at PRJNA1082657. Redkmer Pipeline was executed using CQ threshold values suggested by Papathanos and Windbichler (Xmin of 1.5, Xmax of 2.5, and Ymax of 0.3)(46). No modifications were applied to RedKmer configuration files. Steps 7 (generation of sex chromosome-specific k-mers) and 8 (off-target analyses) of the RedKmer pipeline were modified to generate a list of candidate Y-specific k-mers from the long-read nanopore Y bin. BLASTn tool was used to characterise candidate Y k-mers generated from RedKmer pipeline against all ‘*Anopheles gambiae’* (Taxid: 7165) sequences available in the Gene Bank database (84). BlastN was performed using the following set-up: perc_identity: 100 % and word_size = 25. BlastN results together with information of k-mer abundance were plotted using matplotlib 3.8.0 in Python (85). 25 bps candidate Y k-mers were scanned for the presence of PAM motif at their 5’ or 3’ end, considering a minimum sequence length ≥ 20 bps. The filtering was performed using a homemade Python script. List of candidate Y k-mers can be found in Supplementary File S1.

### 3 Cytological characterization of candidate Y-k-mers

DNA FISH was used to cytogenetically validate the Y-specificity of the target site selected (gRNA^AgY53B^, gRNA^AgY53A^). To generate probe sequences for in situ hybridization, the target site sequence was aligned against the Nanopore Y-bin using bowtie1 and reads showing 100 % matching were selected and included in a separate sub-bin containing only long-reads with perfect matches with the target site. Then, the kmer-specific sub-bins were aligned using Clustal-Omega and the consensus sequence was visualized using Jalview (86,87). A region of 44 bp that includes the 25 bp target site sequence was selected as a reference for the generation of the fluorescent probes. 3’ Cy-3 labelled oligo corresponding to the reference sequence was synthesized by Eurofins and was used as a probe during FISH experiments. Probe specific for the X-linked rDNA locus was generated by PCR as shown in Timoshevskiy et al. 2012 (88). Male mitotic chromosome preparations from WT *An. gambiae* G3 mosquitoes and DNA FISH. were performed according to previous studies (15,88). Briefly, chromosome preparations were treated with Sodium Saline Citrate (SSC) solutions, digested using trypsin solution and fixed using formaldehyde (4 %). Next, chromosome spreads and probes dissolved in the hybridization buffer were denatured together for 4.30 min at 68 °C and hybridized at 37 °C overnight. Subsequently, after the washing steps, the microscope slides were mounted with Prolong Antifade + DAPI, sealed with rubber cement (Cytobond) and stored at Room Temperature (RT) for at least 2 h before confocal analysis. List of containing oligonucleotide sequences and primers used to generate the probes can be found in Supplementary File S2.

### 4 Generation of CRISPR^AgY53B^ construct

The p167 vector, containing attp, piggyBac recombination sequences, 3xP3::eRFP fluorescent marker and βtub::Cas9 (33), was used for Golden Gate cloning. BsaI was used in the reaction to insert the kmer-AgY53B gRNA sequence GAATAGAATCAGAAAAGT. Primers kmer-AgY53B-F: 5’-TGCTGAATAGAATCAGAAAAGT-3’ and kmer-AgY53B-R: 5’-AAACACTTTTCTGATTCTATTC-3’, were used during oligo-annealing to obtain the final transformation construct p16710 (CRISPR^AgY53B^). To verify the insertion of the gRNA the final construct was sequenced via Eurofins. Primer sequences can be found in Supplementary File S2. Plasmid annotated sequence can be found in Supplementary File S7.

### 5 Generation and characterization of transgenic strains

Embryos from *An. gambiae* G3 strain (referred to as wild-type, WT) were injected with a mixture of 200 ng/µl of CRISPR^AgY53B^ and 200 ng/µl of vasa-driven piggyBac transposase helper plasmid (83) using a Narishige IM-400 microinjector and a Narishige MM-94 micromanipulator mounted on a Nikon Ts2R inverted microscope. Two days following the injection, larvae reporting transient expression of RFP marker were isolated using an Olympus MVX10 stereo microscope. Adult mosquitoes expressing RFP were crossed individually to WT to verify the transformation of the germline and obtain the transgenic lines. Based on the RFP inheritance pattern detected in the progeny, single insertion events were selected and inverse PCR was used to characterize the genomic location of the insertion site for two transgenic lines. Inverse PCR was performed as previously described (32,33) and detailed information about the results can be found in Supplementary File S3. The transgenic strains generated were maintained by backcrossing transgenic females to WT males each generation.

### 6 Genetic crosses and phenotypic assays of AgY53B strains

HRs and adult sex ratio were assessed by crossing (40 to 50) two-three day-old adult male mosquitoes heterozygous for CRISPR^AgY53B^ transgene to an equal number of WT *An. gambiae* for 4 days. As a control, negative siblings males were crossed to an equal number of WT *An. gambiae* females. This set-up was used for both the strains harbouring CRISPR^AgY53B^ transgene, AgY53B/2R and AgY53B/2L. On the fifth day, females were blood-fed using Hemotek with cow blood. 2 days after the blood-feeding females were placed individually into a 300 ml beaker filled with water and lined with filter paper and given up to three days to oviposit. For each individual, the number of eggs laid as well as the larvae hatching were counted. When no larvae were observed in the progeny, the spermathecae of the female were dissected to verify the presence of sperm. Females with no sperm in their spermathecae, which laid eggs that did not hatch were removed from the hatching rate counts. Females that did not lay eggs but were inseminated by the males were considered to have a hatching rate value of 0. Larvae were reared until adulthood and the sex ratio was scored for each progeny (Supplementary File S4). Statistical tests were performed using GraphPad Prism version 10.0.0 for Windows, GraphPad Software, Boston, Massachusetts USA.

### 7 Genetic crosses and phenotypic assays on Y-shredder strains in combination with X-shredder and Vasa2:Cas9 strains

To obtain Ag(PMB)1-AgY53B/2L or 2R trans-heterozygous individuals, heterozygous Ag(PMB)1 males were crossed *en masse* with AgY53B/2L or AgY53B/2R females. From the offspring of the cross, L1 larvae were sorted using a ‘COPAS Mosquito’ large particle flow cytometer (Union Biometrica) using a 488-nm solid-state laser and emission filter for the detection of DsReD. Trans-heterozygous larvae were selected and separated according to the intensity of RFP fluorescence. 30 trans-heterozygous males of each transgene combination were crossed to 30 WT *An. gambiae* females and allowed to mate for 4 days before blood feeding. The number of eggs laid, the larval hatching rate and the adult sex ratio were measured as previously described (Figure 5, Supplementary File S5). Further, additional trans-heterozygous males generated from the parental crosses were isolated to perform DNA FISH. To obtain trans-heterozygous Vasa2:Cas9-AgY53B/2R females, 50 Vasa2:Cas9 heterozygous females were crossed to 50 heterozygous AgY53B/2R males allowing them to mate for 5 days *en masse*. Females were blood-fed on day 6^th^ and given up to 3 days to oviposit eggs into a cup. The progeny of the cross was sexed and screened for the presence of YFP (Vasa2:Cas9) and RFP (AgY53B/2R) using Olympus MVX10 stereo microscopes. The progeny of this cross was highly female-biased due to the presence of the AgY53B/2R construct, active during male spermatogenesis. Sibling females harbouring only RFP fluorescent marker (AgY53B/2R) were isolated and used as control (Figure 6). On the contrary, to obtain trans-heterozygous Vasa2:Cas9-AgY53B/2R males, 50 Vasa2:Cas9 heterozygous males were crossed to 50 heterozygous AgY53B/2R females. The progeny of the cross was sexed and screened as previously described and 50 trans-heterozygous males were isolated. As a control, 50 heterozygous AgY53B/2R siblings males were selected from the progeny. Trans-heterozygous and heterozygous (control) adult mosquitoes from both crosses were mated with an equal number of WT *An. gambiae* of the opposite sex. For each cross, 30 females were allowed to lay individually 3 days post blood-feeding. The number of eggs laid, the larval hatching rate and the adult sex ratio were measured as previously described (Supplementary File S5). For these last crosses, the adult sex ratio was recorded only from 15 progeny randomly selected from trans-heterozygous progeny. For the controls, 5 progeny were selected. Further, additional trans-heterozygous males were isolated to perform whole testis DAPI staining and DNA FISH. Statistical tests were performed using GraphPad Prism version 10.0.0 for Windows, GraphPad Software, Boston, Massachusetts USA.

### 8 Whole testis DAPI staining and imaging

At least 5 testes were dissected in 1X PBS from 1-2 days old adult mosquitoes for each genotype under investigation (Supplementary Figure S5). Immediately after dissection testes were transferred into an embryo dish containing 4 % formaldehyde in 1X PBS and incubated for 20 minutes at RT. Then, testes were washed 3 times in a solution containing 0.1 % Tween PBS 1X (PBST) for 15 minutes at RT. Testis were mounted in clean microscope slides with 15 μl of Prolong Antifade gold + DAPI, sealed with Cytobond rubber cement and stored at RT for at least 2 hours before confocal analyses.

### 9 DNA FISH on pupae, adult testis and spermathecae

DNA FISH on strain AgY53B/2L, AgY53B/2R, Ag(PMB)1, Ag(PMB)1-AgY53B/2L, Ag(PMB)1-AgY53B/2R was performed on at least 5 testes dissected from different individuals at late pupal stage as described in method section 1.3. DNA FISH on strain Vasa2:Cas9-AgY53B/2R was performed on testes dissected from 1-2 day-old adults. In this case, to label the X chromosome, an oligonucleotide probe specific to conting_240 was used (15). DNA FISH on female spermathecae was performed according to Haghighat-Khah et al., 2020 (42). Briefly, 30 WT females were crossed with males of the genotype of interest (AgY53B/2L, AgY53B/2R, Ag(PMB)1-AgY53B/2R) and after 6 days females’ spermathecae were dissected in 1X PBS. Each spermatheca was transferred in a clean microscope slide with a fresh drop of 1X PBS and a coverslip was mounted on the drop. A microscope equipped with a phase-contrast filter was used to verify the presence of sperm inside the spermathecae and a flat rubber of a pencil was used to crash the spermathecae allowing sperm to be released. Then, slides were dipped in liquid nitrogen and dehydrated in a series of 50 % (prechilled at -20, for at least 2 hours), 70 % (5 minutes, RT), 90 % (5 minutes, RT) and 100 % (5 minutes, RT) ethanol. After the last dehydration step, slides were let air dry at RT for 30 minutes and stored at RT for up to 1 month. Slides can also be stored at - 20 if not processed within a month. DNA FISH was performed as previously described using probes specific to the X chromosome (rDNA or conting_240 oligo probe) or the Y chromosome(AgY53B oligo probe).

### 10 Microscopy and image analyses

Images were acquired using either Leica SP8 inverted or Leica SP8-STELLARIS 5 Inverted confocal microscopes at Imperial College London FILM facility. Whole testis DAPI staining were visualized using a 40x oil immersion objective while chromosome preparations were observed with a 63x oil immersion objective. Images were analysed using FIJI and Photoshop v2023. In some cases, chromosome pictures were deconvoluted using either Huygens Software 22.10 or the lightning algorithm available in the Leica SP8-stellaris software. A 3D model of mitotic chromosomes was generated from confocal Z-stacks using Huygens Software 22.10 (Supplementary Video, S1). Sperm sex chromosome genotyping on testes and spermathecae was performed using the ImageJ ‘Colocalization Image Creator’ Plugin (53). The software was used to assign sperm to 4 different categories: X-bearing, Y-bearing, XY NDJ and nSC (no sex chromosome) based on the presence of fluorescent signal from sex-chromosome specific probes. Sperm counts and statistics can be found in Supplementary File S6.

## Acknowledgements

We thank Matthew Gribble and Giulia Morselli for experimental assistance and Miles Thorburn, Barbara Fasulo, Brandt Warecki, Alekos Simoni, Priscilla Bascunan, Silke Fuchs and John Connolly for useful discussion. This work was supported by a grant from the Bill & Melinda Gates Foundation and Open Philanthropy. We thank the Facility for Imaging by Light Microscopy (FILM) at Imperial College London for the microscopy analysis.

## Author contributions

M.V, N.K, F.B designed the research, and M.V, N.K and T.C performed bioinformatic analyses. M.V, J.L, K.K, T.C, L.M, S.G performed the research. M.V, N.K and F.B analysed the data. M.V, F.B, J.L and N.C wrote the paper with the input from A.C.

## Supporting Information

**S1 Video: 3D model showing spatial distribution of Y specific fluorescent probes.** 3D model generated from confocal Z-stacks using Huygens software. Yellow shows the signal from the probe specific to gRNA AgY53A, while light blue shows the signal from gRNA AgY53B. Autosomes and X chromosome are shown as red background in 2D for spatial reference. AgY53A signal is confined to the distal end of the Y-chromosome. An additional spot located at around half of the chromosomal length is also detectable in the 3D model. Similarly, AgY53B signal is located at the distal end of the chromosome, however, it can also be detected across different ‘spots’ along the entire chromosomal length. The signals from the target sites specific probes are partially overlapping at the distal end of the Y chromosome.

**S1 File: List of candidate Y k-mers.**

**S2 File: DNA FISH probe sequences.**

**S3 File: Inverse PCR primers and plasmid flanking sequences.**

**S4 File: Phenotypic assay data for strains AgY53B/2L and AgY53B/2R.**

**S5 File: Phenotypic assay data for trans heterozygous strain Ag(PMB)1-AgY53B/2L, Ag(PMB)1-AgY53B/2R, Vasa:Cas9-AgY53B/2R.**

**S6 File: Sperm sex chromosome genotyping raw data.**

**S7 File: CRISPR^AgY53B^ plasmid sequence.**

**S1 Figure:**
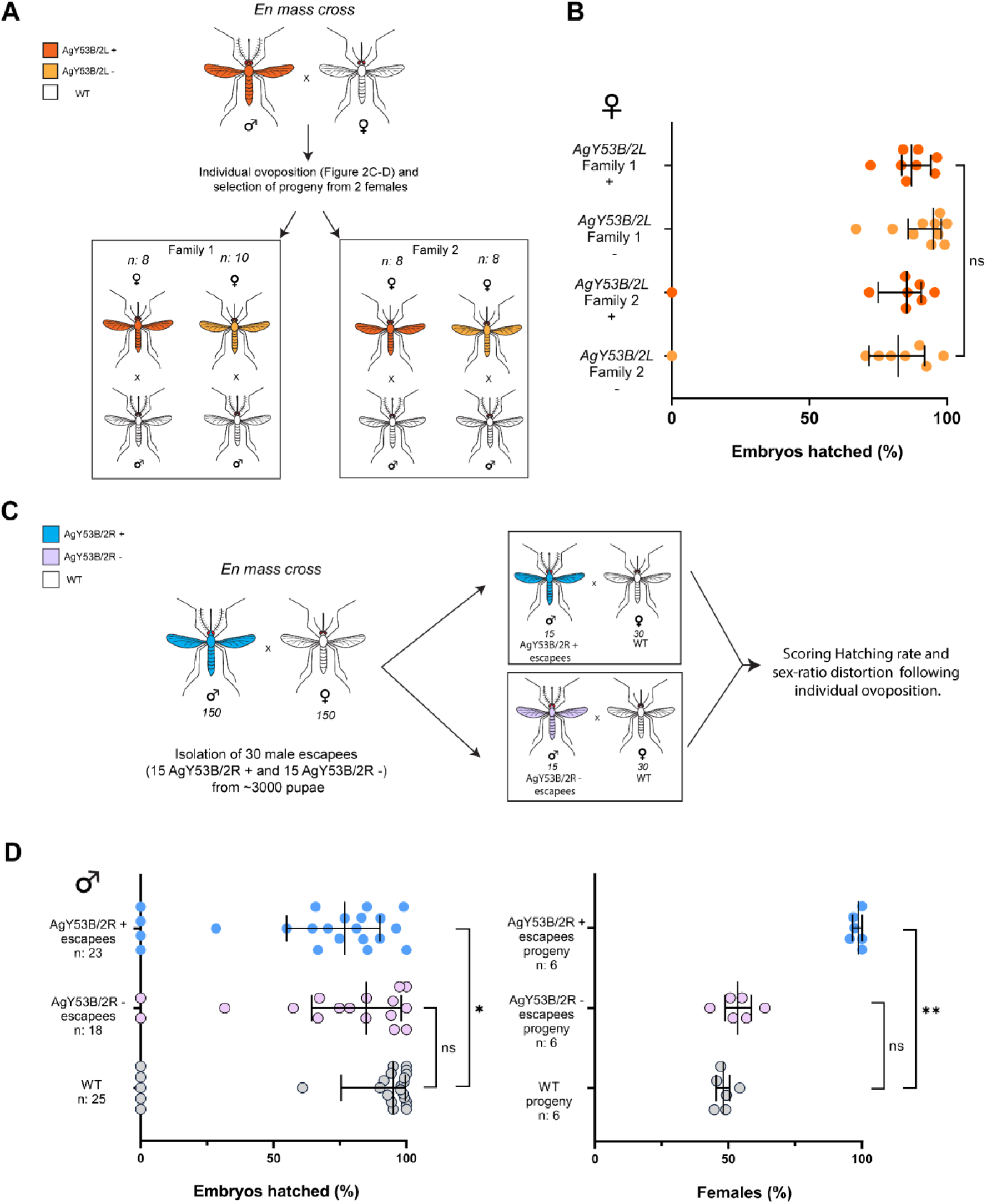
Phenotypic assays on progenies generated from SSD strains. (A) Schematic of the genetic cross used to isolate two different families in the progeny of AgY53B/2L + males, this refers to the phenotypic assay shown in Figure 2-C (main text). Transgenic and non-transgenic female progeny (AgY53B/2L + and AgY53B/2L -, correspondingly) were separated in different cages and crossed with 20 WT *An. gambiae* males. Number of females used for each cross are shown. Following blood feeding, females were allowed to lay individually and the HR was scored (B). No significant reduction in the HR was detected in AgY53B/2L + as well as in non-transgenic sibling females (Dunn’s multiple comparison test, P value > 0,05). (C) Schematic of the genetic cross used to isolate escapee males from strain AgY53B/2R. Transgenic and non-transgenic escapee males were crossed to 30 WT females in two separate cages and a phenotypic assay was performed as previously described. In addition, sex ratio for six randomly selected progenies was analysed (D). A slightly, but significant, reduction in the HR (Dunn’s multiple comparison test, *P value* = 0.0448) was observed when comparing AgY53B/2R + escapee males with the WT control. The analyses of the progenies randomly selected shows high female bias for AgY53B/2R + males and value in line with Mendelian segregation for AgY53B/2R – and WT controls. Median and interquartile ranges are shown.

**S2 Figure:**
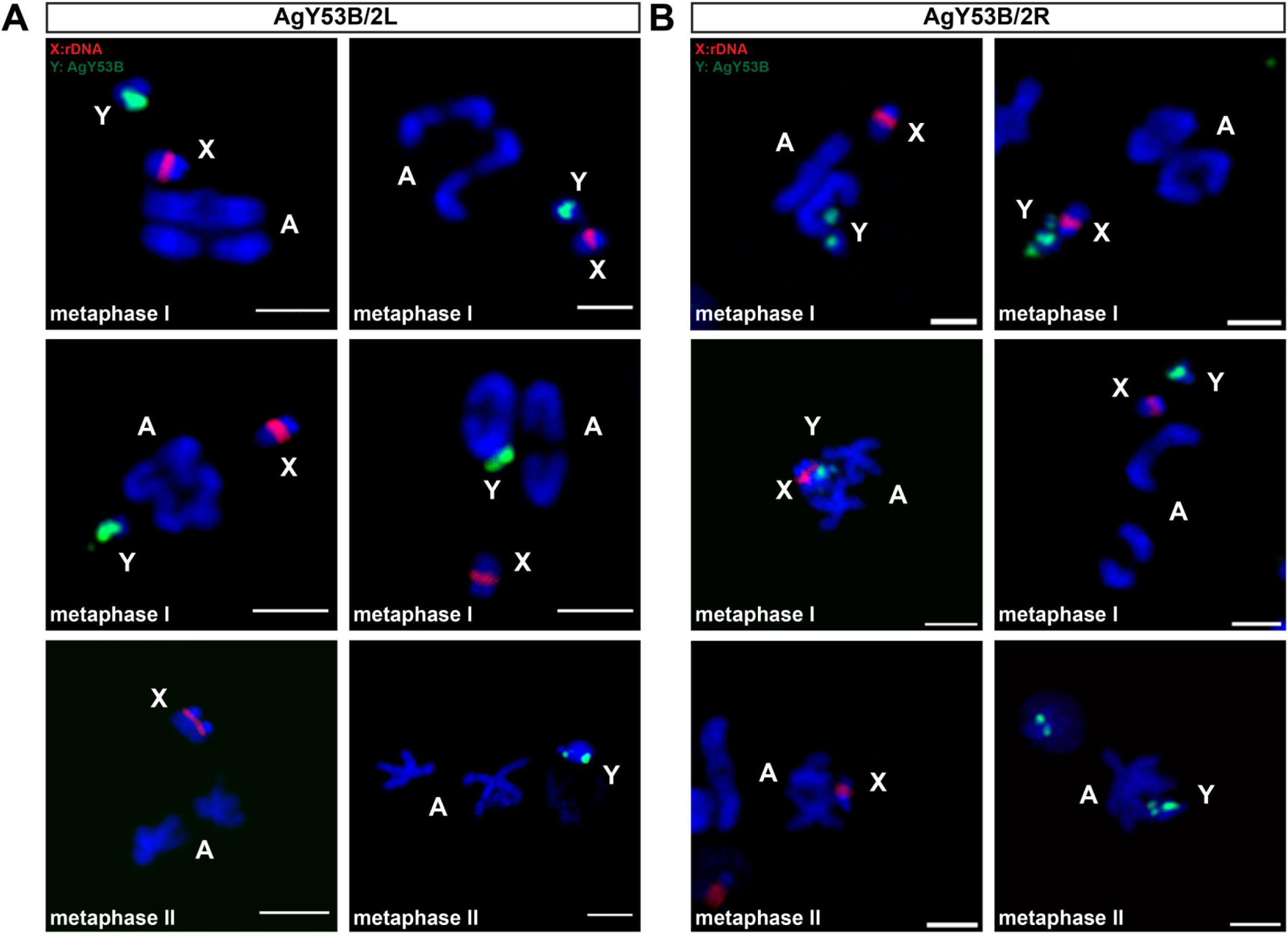
Metaphase chromosomes obtained from sex distorter strains shows no evident alterations. (A) Both sex chromosomes are detectable in metaphase I of cells obtained from the testis of AgY53B/2L strain. Chromosomes in metaphase plate and chiasmata can also be detected. Cells in Metaphase II bearing X or Y chromosome can also be observed indicating that correct segregation of sex chromosome can still occur despite Y-chromosome lagging. (B) A similar scenario can also be observed in the strain AgY53B/2R where metaphase I and II cells show the presence of both sex chromosomes. For comparison with metaphase karyotype from WT strain see Liang and Sharakhov, 2019 (15). Scale Bar = 3 um. Blue = DAPI, Red = X chromosome-specific probes (X: rDNA), Green Y chromosome-specific probe (Y: AgY53B).

**S3 Figure:**
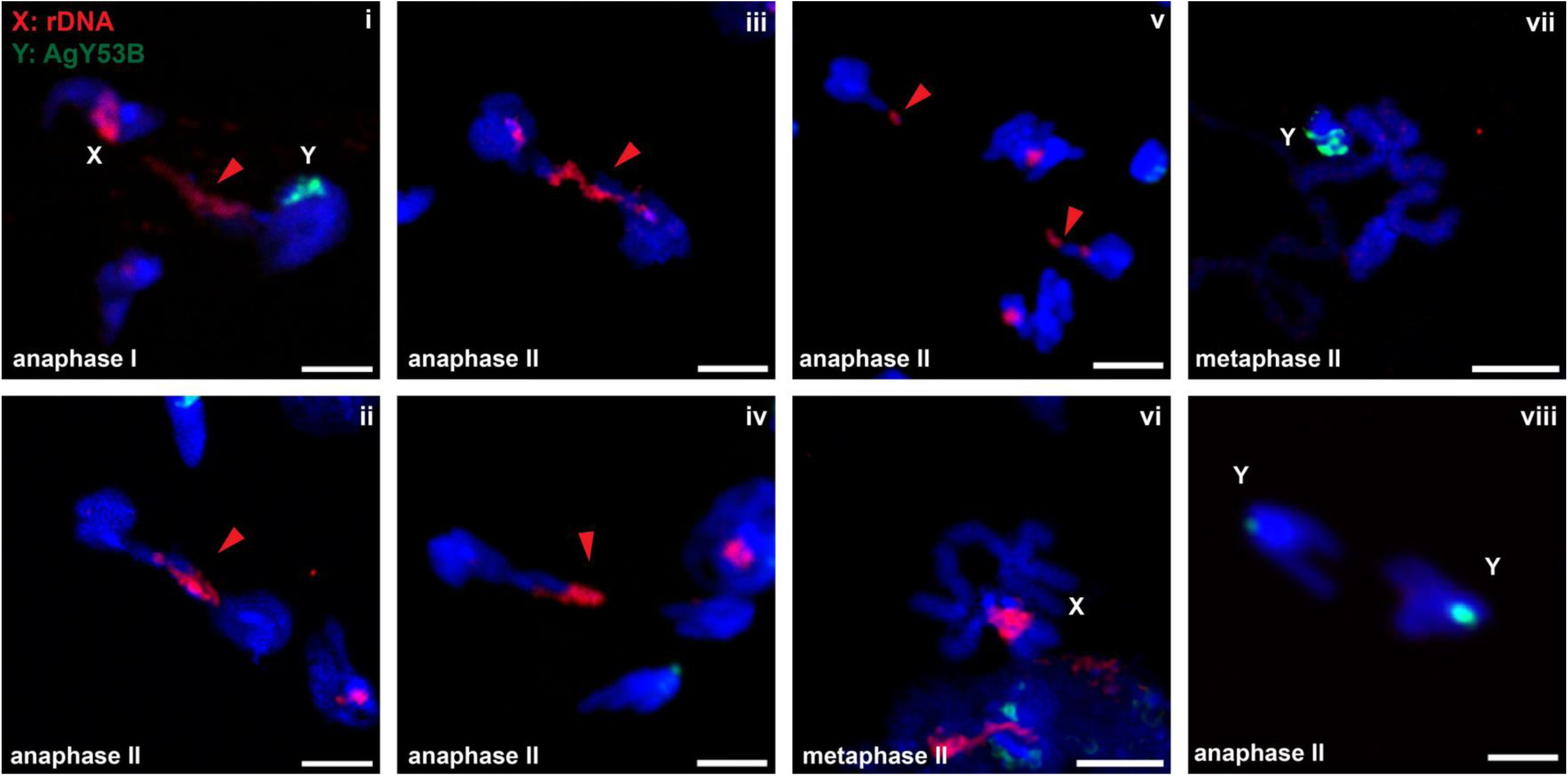
Anaphase bridging/lagging of the X chromosome can be observed in strain Ag(PMB)1 following shredding at the X-linked rDNA locus. DNA FISH on meiotic chromosomes obtained from Ag(PMB)1 testis. Panel i shows X-chromosome lagging during anaphase I, while Y chromosome segregate at the pole of the cell normally. Panels ii-iii-iv-v show X-chromosome lagging during anaphase II. Panel vi and vii show sex chromosomes metaphase II. Panel viii shows cells in anaphase II with no lagging of the Y chromosome. Red = X chromosome specific probes (X: rDNA). Green = Y chromosome-specific probe (Y: AgY53B). Scale bar 3 µm. red arrowhead indicates the lagging chromosome.

**S4 Figure:**
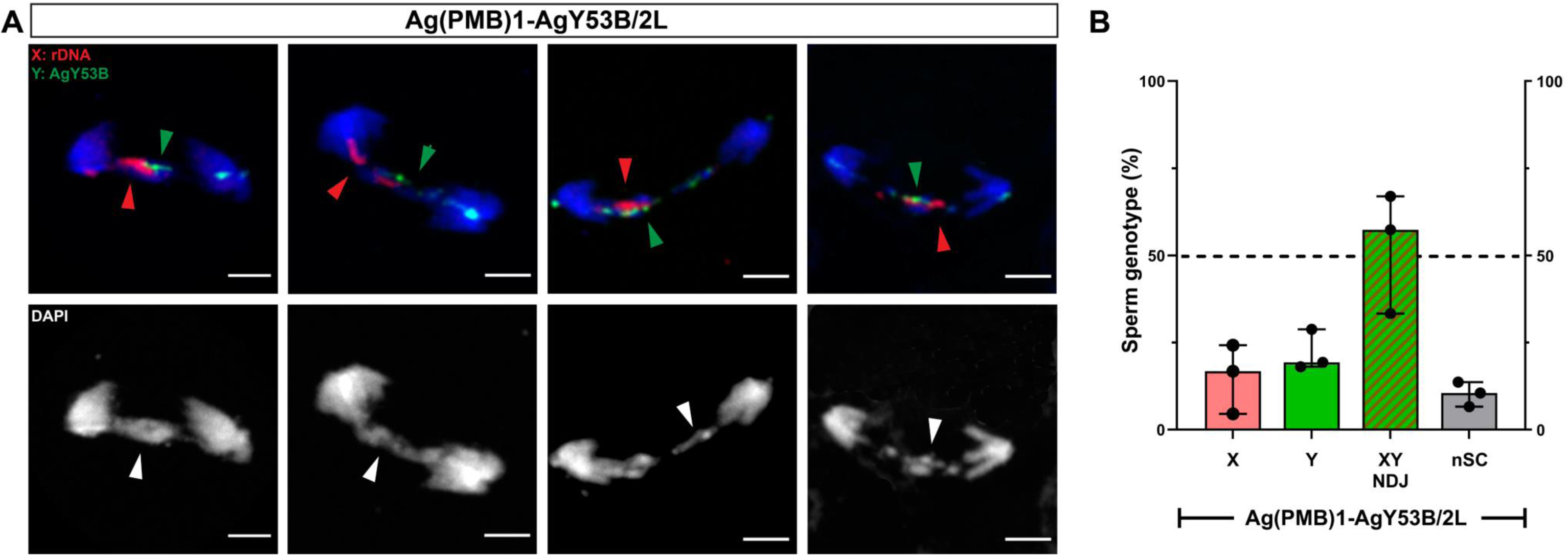
Cytological investigation performed on Ag(PMB)1-AgY53B/2L trans heterozygous male testis. (A) DNA FISH reveal lagging sex chromosomes during meiotic anaphase I in trans heterozygous Ag(PMB)1-AgY53B/2L males. The lagging pattern is similar to the one detected in trans heterozygous Ag(PMB)1-AgY53B/2R shown in Figure 5-C (main text). Red and green arrowheads indicate lagging sex chromosomes. Red = X chromosome-specific probes (X: rDNA). Green = Y chromosome-specific probe (Y: AgY53B). Scale bars = 3 µm. (B) Sperm sex chromosome genotypes of testes obtained from 3 trans heterozygous Ag(PMB)1-AgY53B/2L males. Median of the percentage of X-bearing = 16.75 %, Y-bearing = 19.29 %, XY-NDJ = 57.36 %, nSC = 10.53 %. We observed a higher proportion of XY NDJ in testis dissected from Ag(PMB)1-AgY53B/2L compared to Ag(PMB)1-AgY53B/2R males (see Figure 5D, median XY NDJ of the strain Ag(PMB)1-AgY53B/2R = 19.65 %). Total number of sperm counts: X = 55, Y = 81, XY NDJ = 224, nSC = 36. In each bar median with Standard Deviation (SD) is shown.

**S5 Figure:**
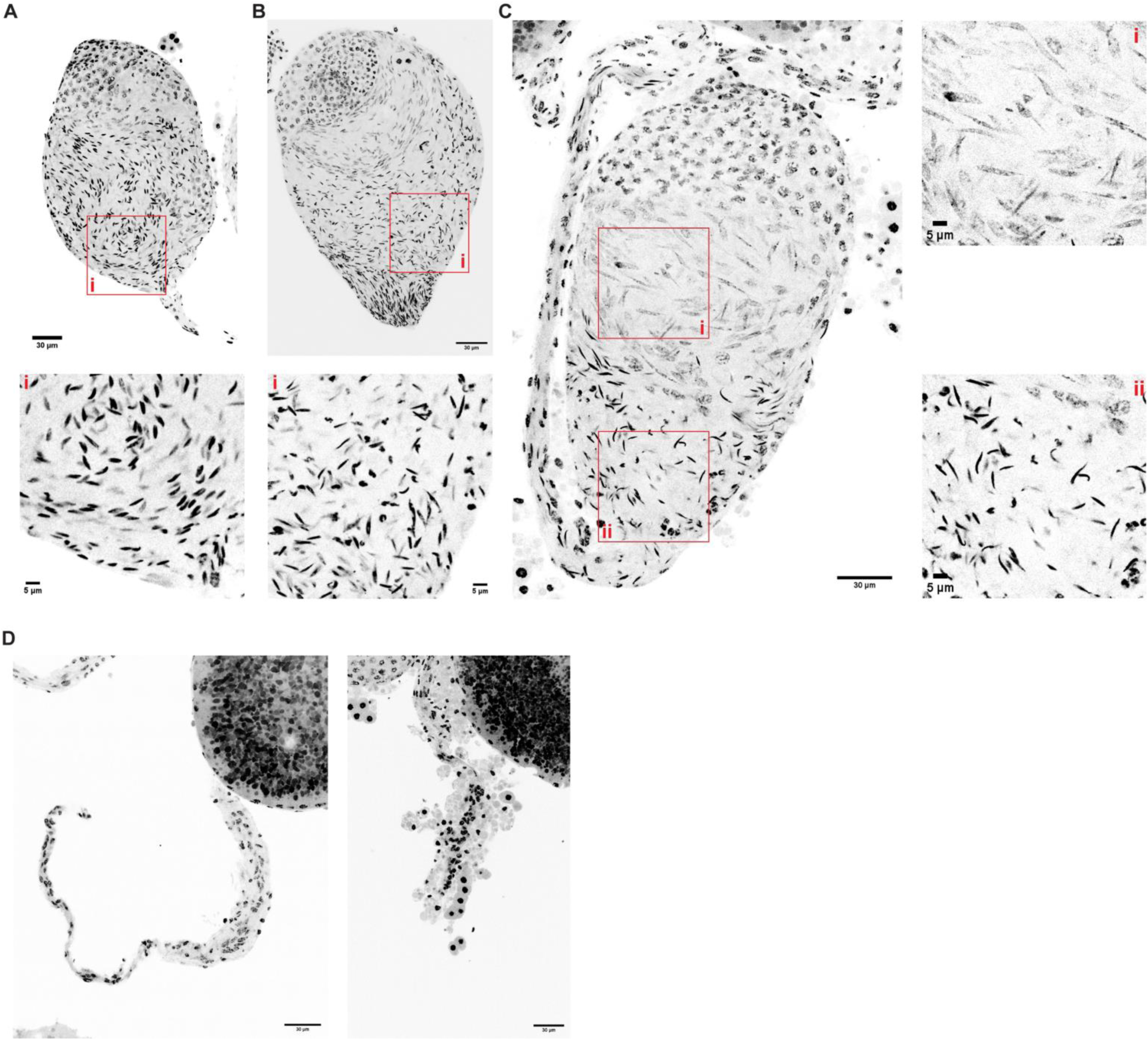
Whole organ DAPI staining reveals details of testes phenotype and sperm defects in trans heterozygous Vasa2:Cas9-AgY53B/2R males. (A) Control sibling males harbouring no genetic construct. We observed sperm nuclei showing normal arrow-like shape (B) Testis dissected from sibling males harbouring only CRISPR^AgY53B^ construct (AgY53B/2R). In these males some sperm show chromatin condensation defects. This can be linked to the presence of defective sperm with X-Y NDJ according to the previous analyses on sperm sex chromosome genotype. (C) Testis dissected from trans heterozygous Vasa2:Cas9-AgY53B/2R males. Spermatids and mature sperm show evident chromatin condensation defects, with elongated and irregular shapes (C-i-ii). Nevertheless, some sperm show a normal-like shape (C-ii). (D) Atrophic-like testes dissected from trans heterozygous Vasa2:Cas9-AgY53B/2R males. In these testes no mature sperm were detected. These testes have a smaller size and contains a lower number of cells if compared to the testes in A, B and C. All the testes samples were dissected from one days old adult males. At least 4 testes for each group were analysed.

**S6 Figure:**
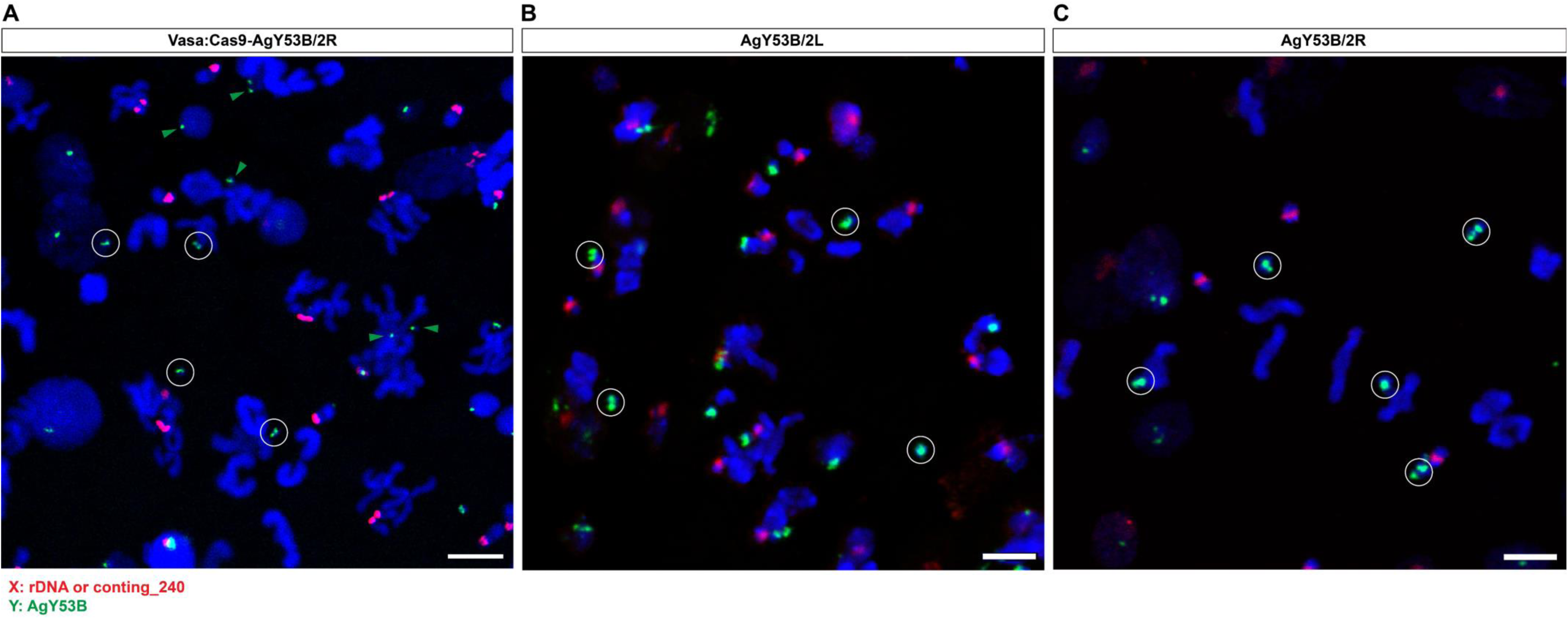
DNA FISH on meiotic chromosome spreads obtained from Vasa:Cas9-AgY53/2R trans heterozygous males (A), AgY53B/2L (B) and AgY53B/2R (C) strains. (A) Pre-meiotic shredding of the Y chromosomes is achieved using Vasa promoter in combination with CRISPR^AgY53B^ construct present in the strain AgY53B/2R. Signals from target site specific probe reveal the presence of a small and fragmented Y chromosomes distributed across the meiotic chromosome spread. The size of the signal from the probe specific to the target site AgY53B is smaller when compared to probe signal in B and C (white circles). White circles highlight the difference in the sizes of the Y chromosomes between the 3 strains. White circles diameter is 2.5 µm. Green arrowheads indicate fragmented Y chromosome. As shown in Figure S3, signal from AgY53B probes cover most of the Y chromosome in prophase/metaphase cells. Red = X chromosome specific probes, rDNA locus or conting_240. Green = Y chromosome specific probe, AgY53B. Scale bars 5 µm.

